# Seasonal hepatic plasticity follows a temporal response hierarchy in a Neotropical frog

**DOI:** 10.64898/2026.07.26.740824

**Authors:** Lilian Franco-Belussi, Luciana T. Moraes, Classius de Oliveira, Carlos E. Fernandes, Diogo B. Provete

**Author notes:** These authors contributed equally to this article (Lilian Franco-Belussi, Luciana T. Moraes).

## Abstract

**Background:** Seasonal climate variation drives physiological change across levels of biological organization in ectotherms, yet whether these responses follow a predictable temporal hierarchy remains untested in wild populations. In a year-round field study of the Lesser Treefrog (*Dendropsophus minutus*, Anura: Hylidae) in São José do Rio Preto, southeastern Brazil, we sampled 40 to 68 adult males across a full annual cycle to quantify seasonal variation in four hepatic phenotypic modules spanning multiple organizational levels: liver histochemistry (pigments and glycogen; intracellular), cell and nucleus morphometry (cellular), tissue volumetric composition of hepatocytes, sinusoids, melanomacrophage centres and portal structures (tissue), and whole-body somatic indices of liver mass and body condition (organismal). We asked whether lower organizational levels respond faster to seasonal climate, as predicted by the bottom-up cascade framework originally proposed for toxicant exposure, or whether alternative hierarchies emerge when climate acts first through whole-organism physiology. To test whether climatic change precedes each biological response, we extended Procrustean superimposition to incorporate time lags, tested both forward and reverse directions, and complemented it with phenotypic trajectory analysis, phase analysis of monthly change, and lagged regression against temperature, precipitation, humidity and drought.

**Results:** Somatic indices and tissue composition tracked climate with no detectable delay, whereas histochemistry and cell morphometry accumulated a three-month delay, and tissue composition in turn led somatic indices by one month. Trajectory analysis separated the modules into a fast-responding group (somatic indices, histochemistry and morphometry) and a slower tissue-composition tier, a two-tier structure confirmed by paired bootstrap resampling in which the slow tier differed from every fast module in both displacement rate and total path length. The four modules were largely independent through time, and the two strongest concordances linked climate to tissue composition and to somatic indices.

**Conclusions:** These results reveal a temporal response hierarchy in which seasonal climate acts first on whole-organism energy balance and then propagates to tissue architecture and intracellular processes with module-specific delays, showing that the direction of the hierarchy depends on the nature of the environmental driver. We also introduce a general analytical framework for testing temporal precedence in short multivariate biological time series.

## Background

Interpreting temporal variation in morphological and physiological traits requires a framework grounded in evolutionary theory. Phenotypic plasticity, the ability of a genotype to produce different phenotypes in response to environmental cues, is a primary mechanism for coping with temporal environmental heterogeneity [1, 2]. Understanding how fast different phenotypic components respond to environmental change, and whether these responses follow a predictable sequence, is central to predicting organismal performance under fluctuating conditions [3, 4]. Theory predicts that strong plastic responses should evolve in environments with low variability [5], whereas unpredictable environments select for reduced plasticity to avoid the fitness costs of maladaptive mismatches between phenotype and the environment (reviewed in [4]). This theoretical framework has been tested in short-lived, single-celled organisms [6]. However, little is known on how the phenotype of long-lived vertebrates with complex life cycles, such as frogs, respond to mid- and long-term environmental changes. Additionally, few studies have estimated the speed at which the phenotype changes through time [3], focusing on comparisons between populations at a given time point. Nonetheless, this knowledge gap hinders our understanding of how organisms might respond to climate change [2] and if their phenotype can cope with the predicted rate of environmental change.

Frogs are excellent models to investigate temporal phenotypic plasticity because, as ectotherms with highly permeable skin, their activity, physiology, and energy balance are tightly coupled with external environmental conditions [7]. This tight coupling means that seasonal climate variation should produce measurable, hierarchically organized responses across different levels of biological organization, from intracellular biochemistry to whole-organism energy allocation (see also [8]). While much research has focused on developmental plasticity during the short-lived aquatic larval stage (e.g., [9]) or on how these early environments carry-over to adult morphology [8], adult frogs can live for several years in highly seasonal terrestrial habitats. Consequently, adults must continuously adjust their internal morphology and physiology to cope with cyclical, mid- to long-term seasonal variations throughout their lifespan. Furthermore, plastic responses are often aligned with the major axes of standing genetic variation (*g_max_*), giving organisms high evolutionary potential and suggesting that selection on environmentally induced phenotypes can be effective [10, 11]. However, this efficacy depends on phenotypic modularity, i.e., the organization of the phenotype into semi-independent units that can evolve distinctly [12]. Modules exhibiting different speeds of plasticity allow natural selection to refine the environmental sensitivity of certain traits without disrupting the functional stability of others, a fundamental feature of evolvability [13]. Thus, by characterizing the temporal trajectory of distinct hepatic modules in response to environmental fluctuations, this study explores how the hierarchical architecture of the frog liver, from intracellular histochemistry to whole-organism somatic indices, reflects an adaptive strategy that might fuel future evolutionary change and species persistence under shifting climates.

Previous studies have analyzed how the phenotype of frogs changes seasonally, such as body size (e.g., [14, 15, 16]), testis size, and gonadal development (e.g., [17, 18]), body condition (e.g., [19]), and liver mass [20, 21]. Liver tissue characteristics are also known to change seasonally in response to environmental factors, with glycogen, body fat, and blood glucose increasing at the onset of the breeding season [22, 23], and liver histology differing markedly between pre- and post-reproductive periods [24], throughout which frogs deplete their energy reserves. These seasonal changes in substances produced by the liver are accompanied by changes in hepatocytes [25]. However, these studies typically analyzed one or a few of those traits in isolation, without explicitly testing whether different levels of biological organization respond at different speeds to the same environmental cue. However, changes in hepatic characteristics occur in synchrony, as substances produced by hepatocytes require a modification of the cell and nucleus size and the liver tissue itself [26, 27]. Despite all of that, no study has quantified the temporal trajectory of multiple hepatic components jointly, especially in tropical species that do not hibernate (but see [28]). This approach would allow us to understand the chain of causation that leads to changes in the phenotype observed at a given time of the year.

Cell and tissue density, estimated through stereological point counting, provides a tissue-level perspective on liver physiology [24, 29]. The relative proportions of hepatocytes, sinusoids, melanomacrophage centers, and portal structures reflect the functional state of the liver parenchyma at a given time, integrating metabolic activity, vascular remodeling, and immune function [29, 30, 31]. Seasonal changes in these proportions have been documented in temperate frogs [25, 28, 32], but their temporal relationship to other hepatic traits remains unexplored.

At the organismal level, somatic indices, such as the hepatosomatic index (HSI) and the Scaled Mass Index (SMI; [33]) provide integrative measures of energy storage and body condition. These indices respond rapidly to changes in food availability and metabolic demand because they reflect whole-organ mass relative to body size [34, 35]. For example, HSI is related to glycogen and lipid storage, protein synthesis, and detoxification activity [34]. SMI correlates with the total amount of lipids and proteins stored [36] and has been widely used to detect differences in organismal health status associated with environmental change in frogs [19, 37]. Together, tissue density and somatic indices represent the higher end of the biological organization hierarchy, where changes are expected to manifest later and more slowly than at the intracellular level [38]. The circannual rhythm of energy reserves in frog liver — including glycogen, lipids, and pigments — has been well documented in temperate species [22, 23, 39, 40, 41], providing a rich physiological context for interpreting seasonal patterns [37]. However, such patterns have not been fully explored in tropical vertebrates.

The frog liver is an important metabolic organ and responds to several environmental stimuli [34], such as UV [42], temperature (e.g., [43]), and xenobiotics (e.g., [27, 44, 45, 46]). Liver pigments (melanin, hemosiderin, and lipofuscin) reflect oxidative stress and immune function [47, 48], while glycogen content integrates metabolic demands over weeks to months [49, 50]. In year-round reproductively active tropical species, such as *Boana prasina*, these seasonal metabolic changes occur even without hibernation [51], suggesting that seasonal liver plasticity is a general feature of anuran biology, rather than exclusively linked to dormancy cycles. However, previous studies that evaluated the relationship between xenobiotics and hepatocyte morphometry (e.g., [27]) were mostly experimental and ran from a few days to weeks. Additionally, these studies often used tadpoles (e.g., [27]), which live for 40 to 90 days ([52] and references cited therein). Thus, the duration of previous experimental studies covered a small fraction of the organismal life span. Adult, small-sized hylid treefrogs live from 3 to 4 years [53]. Therefore, realistic studies that investigate how hepatic morphological characteristics change through time in response to extraneous factors, like climate, should cover a longer period of the study organism’s life. Beyond hepatic physiology, this might have implications for ecotoxicology, because oft-used short-term laboratory experiments testing the effects of xenobiotics on hepatic traits might be overestimating the impact of these substances, while neglecting the adaptive capacity of organisms.

The liver is also particularly well suited to a multi-level analysis of temporal plasticity. As the central metabolic organ of ectotherms, it integrates energy storage, detoxification, and immune function [34], and its energy reserves show well-documented circannual rhythms in frogs [22, 41, 51]. Intracellular pigments and metabolites, hepatocyte dimensions, tissue composition, and organ mass can all be quantified from the same histological sections and biometric measurements, giving simultaneous access to four organizational levels within a single tissue.

Complex phenotypes are organized into semi-independent modules, i.e., sets of traits that covary strongly among themselves, but weakly with traits in other modules [12]. Modularity allows different aspects of the phenotype to respond independently to selection or environmental variation, which facilitate evolvability by allowing selection to modify one set of traits without disrupting others [13]. Whether the temporal dynamics of phenotypic modules also exhibit hierarchical structure, with some modules responding faster than others to environmental change, is a question that connects modularity theory to the study of phenotypic plasticity. In the context of hepatic physiology, the four phenotypic modules we examine: intracellular metabolites (histochemistry), cell dimensions (morphometry), tissue composition (volumetric density), and whole-organism condition (somatic indices) correspond to distinct functional and structural levels of the liver. If these modules are genuinely semi-independent, they may respond to seasonal environmental variation at different speeds and with different temporal lags. Bayne et al. [38] formalized this intuition for toxicant exposure, proposing that perturbations enter the organism at the molecular level and cascade upward through cellular, tissue, and organismal levels. However, whether this temporal hierarchy holds under natural, cyclic environmental variation, in which the environmental signal enters through ecological pathways (food, water, temperature) rather than through direct molecular disruption, remains untested.

Climate influences hepatic characteristics through multiple pathways that operate at different timescales. Temperature directly affects metabolic rate in ectotherms, altering the rate of glycogen synthesis and breakdown [23], melanin production in melanomacrophage centers [54], and hepatocyte turnover [32]. Rainfall and humidity influence prey availability [55] and reproductive phenology, which in turn determines seasonal cycles of energy intake and subsequent liver storage [28, 51]. UV-B radiation increases melanin area in frog melanomacrophages, presumably as an antioxidant response [42]. At the tissue level, seasonal shifts in metabolic demand can alter the balance between parenchymal and stromal components, as hepatocyte hypertrophy during periods of active storage compresses sinusoidal space and changes the overall tissue architecture [25]. These effects are expected to propagate upward: intracellular changes in metabolite accumulation and cell size should precede tissue-level remodeling, which should in turn precede changes in organ mass and body condition [38]. However, the temporal sequence of these responses has never been explicitly quantified in a wild vertebrate population.

Here, we evaluated how different morphological aspects of the liver vary year-round in an open, natural population of the Lesser Treefrog (*Dendropsophus minutus*). To test this temporal hierarchy (Fig. 1), we used three complementary approaches: (1) Procrustean superimposition (PROTEST) extended for the first time to deal with temporal lags, which tests directional concordance between multivariate datasets at different time delays; (2) phase analysis of monthly displacement speed in ordination space, which identifies when each phenotypic module undergoes its greatest rate of change; and (3) lagged regression of climate on biological principal component scores, which estimates the optimal response delay for each module. We hypothesize that hepatic metabolic responses are more sensitive to temporal variations than cellular, tissue, and organismal attributes (Fig. 1), following Bayne et al. [38]. Therefore, we expect that substances, such as lipofuscin and hemosiderin to change more rapidly than cell size, for example. The hot and rainy season in the Tropics usually provides greater food supply for frogs, as arthropods are more active and abundant during this period [55]. The increase in feeding activities alters both metabolism and energy availability. During the rainy season, we also expect an increase in hepatic glycogen storage [51], cell and nucleus size due to increased protein synthesis in the liver [23].

**Figure 1:**
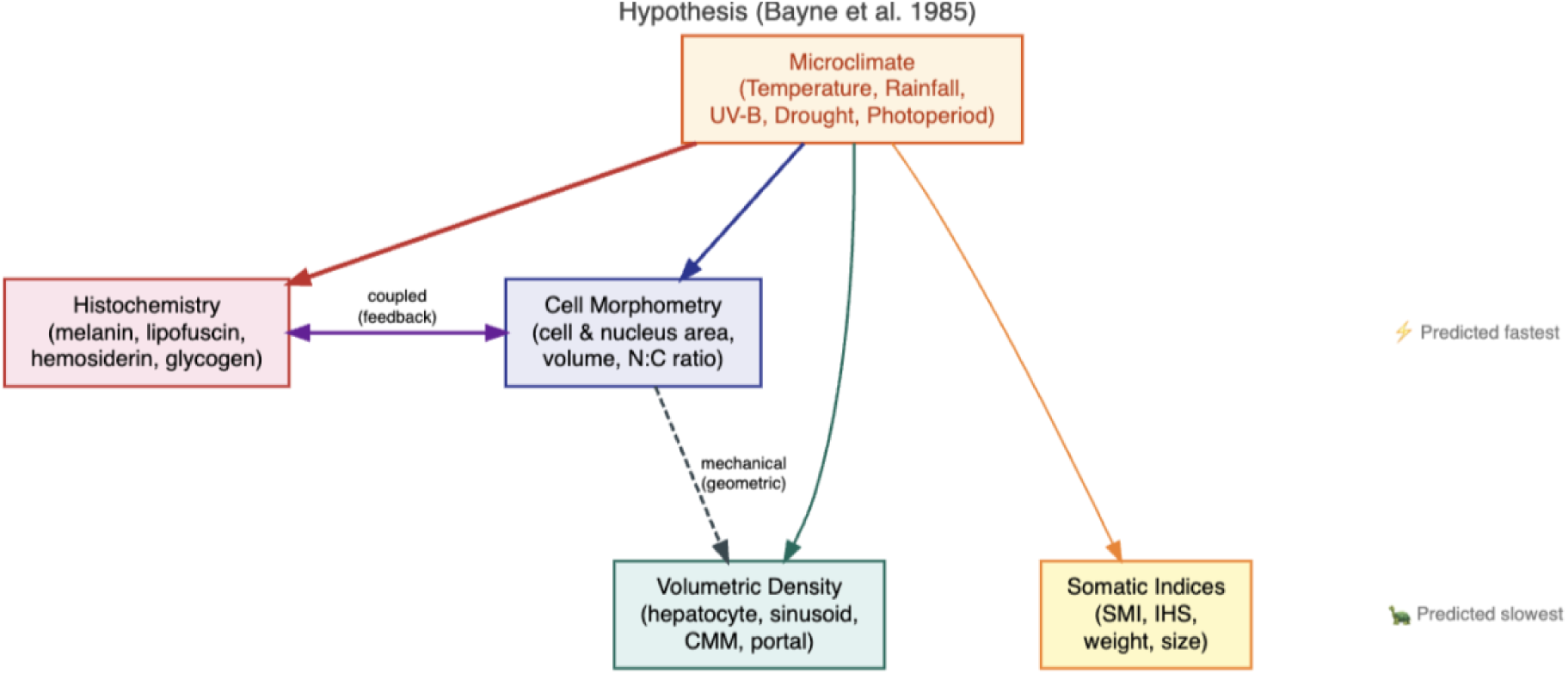
Hypothesized temporal response hierarchy based on Bayne et al.’s (1985) bottom-up cascade framework. Under this prediction, intracellular processes (histochemistry) should respond fastest and organismal traits (somatic indices) slowest. Climate acts on all phenotypic modules (colored arrows). The bidirectional arrow between Histochemistry and Morphometry reflects their positive feedback coupling. The dashed arrow from Morphometry to Density represents the mechanical (geometric) consequence of cell size changes on tissue composition.

## Methods

The following sections follow the ARRIVE guidelines [56] for reporting animal research. We used the MeRIT guidelines [57] to report author contributions in the Methods sections (initials after each task).

### Sampling

**LTM** and **CO** collected 68 adult males of the hylid treefrog *Dendropsophus minutus* from April 2016 to March 2017 (Additional file 1: Fig. S1) by active search at night in temporary water bodies in Engenheiro Schmitt, São José do Rio Preto, São Paulo, southeastern Brazil (49° 21’ 00.8’’ W, 20° 53’ 57.3’’ S). Monthly sample sizes ranged from 4 to 12 individuals (mean = 5). Only adult males were collected, identified by the presence of vocal sacs (minimum SVL = 1.85 cm), to control for the confounding effects of vitellogenesis and egg production in females. Sample size was determined by the number of individuals encountered during active searches and was not predetermined by a power analysis, as this was an observational study. Collection permit was issued by ICMBio (#18573-2). All procedures followed the Brazilian National Committee on Ethics in Animal Research resolution (#34/2017) and the AVMA Guidelines (2020) and were approved by our IACUC (CEUA, #150/2015). Collected specimens are deposited in a zoological collection (DZSJRP).

### Somatic indices

Specimens were euthanized by immersion in a benzocaine solution (5 g/L) by **LTM**. Subsequently, they were dissected to remove the liver. The snout-vent length (cm) was measured with a digital caliper (0.01 mm precision) and body mass (g) was measured with a scale (0.001 g) by **LTM**. From these, **DBP** calculated two somatic indices: the hepatosomatic index (HSI = liver mass / body mass × 100) and the Scaled Mass Index (SMI) following Peig & Green [33], with Standardized Major Axis regression calculated with the R package `smatr` [58]. Higher values indicate better health, greater amount of energy storage, improved changes of surviving harsh environmental conditions, and growth rate [59]. These four variables (SVL, body mass, HSI, SMI) constitute the somatic indices module.

### Routine histological processing

Cross-sections of the whole liver were taken by **LTM**, supervised by **CO**, immediately after euthanasia. The organ was fixed in Karnovsky’s solution, dehydrated in an ethanol series, embedded in historesin (Leica), following Franco Belussi et. al. (2016). Two µm sections of the whole organ made with a rotating microtome (Leica^®^ RM 2265) were used for analysis of structural volumetric density, liver morphometry, quantification of glycogen, and liver pigments (melanin, lipofuscin, and hemosiderin; Additional file 1: Fig. S1). The selection of tissue regions was random and encompassed all hepatic lobes, cortical, and medullary regions.

### Cell and tissue volumetric density

Sections were stained with hematoxylin and eosin (HE). Five images at 400× magnification per specimen were taken by **LTM** of different histological fields, supervised by **CEF**, under a microscope (Leica DM 4000 B) coupled to a digital camera (Leica DFC 280). A 252-point grid was superimposed on each image using ImageJ v. 1.48 [60], and the tissue component underlying each grid intersection was classified into hepatocyte, sinusoid, melanomacrophage center (MMC), blood vessel, or bile duct (Additional file 1: Fig. S1), following Russ & Dehoff [29]. All histological measurements were performed independently by **LTM** and **LFB**, both blind to the collection month of each specimen. No significant discrepancies existed between observers.

### Cell morphometry and stereology

Of the 68 frogs sampled, 40 had complete morphometric data (the same individuals with complete histochemistry); morphometric measurements were unavailable for the remaining 28 individuals as they lacked sufficient material for fine-scale cellular measurements. We obtained 10 images (1,000× magnification) stained in HE in the same equipment described above. We obtained cell-level measurements from the same HE-stained sections used for volumetric density. In each image, **LTM** and **LFB** measured the following variables in 5 hepatocytes chosen at random, totaling 50 measurements per specimen: area (µm^2^), perimeter (µm), major and minor diameter (µm) of the cytoplasm; area (µm^2^), perimeter (µm), major and minor diameter of the nucleus [21]. From these first-order measurements, **LTM** and **LFB** derived two second-order measurements: the nucleus:cytoplasm ratio and cell volume, following Russ & Dehoff [29]. Measurements were taken in Motic 2.0 (Motic Asia, Hong Kong, China).

### Histochemistry

Of the 68 frogs sampled, 40 had complete histochemistry data, with monthly sample size varying from 1 to 7 individuals; hemosiderin, lipofuscin, and glycogen measurements were unavailable for the remaining 28 individuals (Additional file 1: Fig. S1; Additional file 1: Table S2). Glycogen was detected with Periodic Acid-Schiff (PAS) staining by **LTM**. We took five random images at 400x magnification of each specimen. Hemosiderin and lipofuscin detection followed Franco-Belussi et al. [26]. For the analysis of liver pigments, 25 random images of histological fields of each specimen were obtained at 200× magnification. Quantifications of liver pigments and glycogen were done by **LTM** and **LFB** in Image Pro-Plus 6.0 (Media Cybernetics, Rockville, Maryland, USA), following Franco-Belussi et al. [61]. These four variables (melanin area, lipofuscin area, hemosiderin area, glycogen area) constitute the histochemistry module.

### Data Analysis

DBP organized the data into four phenotypic modules (Additional file 1: Fig. S1), each reflecting a different level of biological organization (sensu [38]): (i) histochemistry (intracellular level), (ii) cell morphometry (cellular level), (iii) volumetric density (tissue level), and (iv) somatic indices (organismal level). Representative photomicrographs showing the extremes of each histological module throughout the sampling period are presented in Additional file 1: Figs. S5–S8. These four modules were defined a priori from independent histological protocols and distinct organizational levels, not by any data-driven clustering; accordingly, the Principal Component Analyses described below were applied within each module to summarize its variation, never across modules to impose structure.

Volumetric density variables are proportions estimated by stereological point counting (252-point grid) and are constrained to sum to 100% [62]. Applying standard multivariate methods to such compositional data can produce spurious correlations driven by the closure effect [62, 63]. Therefore, **DBP** applied Compositional Data analysis (CoDa) methods [63, 64] to this module.

First, **DBP** assessed zero prevalence. Blood vessels and bile ducts had high frequencies of structural zeros at the individual level (54% and 50%, respectively) because these portal triad structures are not uniformly distributed across the parenchyma and may be absent from any given set of five microscopic fields. Thus, **DBP** amalgamated blood vessels and bile ducts into a single “portal” component, reducing zero frequency to 38%. Individual-level zeros in the remaining components (hepatocyte: 0%, sinusoid: 0%, MMC: 2.9%, portal: 14.7%) were imputed the count-zero multiplicative (CZM) replacement method [65], appropriate for rounded zeros arising from limited point-counting resolution. Analysis was conducted in the R package `zCompositions` [66].

The imputed four-part composition (hepatocyte, sinusoid, melanomacrophages, portal) was subjected to a centered log-ratio (clr) transformation for subsequent PCA [67, 68]. For volumetric density, the full CoDa pipeline was replicated within each bootstrap iteration to ensure that the bootstrap confidence intervals are computed on the same scale as the observed trajectory metrics.

**DBP** performed separate Principal Component Analyses (PCA) on each phenotypic module and on the climate variables to summarize multivariate variation into orthogonal axes. For the somatic indices, histochemistry and morphometry modules, PCA was computed on the correlation matrix in the R package `FactoMineR` [69] and `factoextra` [70]. For volumetric density, PCA was applied to the clr-transformed data without further scaling [67]. All principal components were retained for each module and computed monthly mean PC scores across individuals. These monthly centroids were then used to trace phenotypic trajectories [71] through the ordination space across the 12 sampling months (April 2016 – March 2017).

**DBP** obtained daily temperature (maximum and minimum), precipitation, relative humidity, and Palmer Drought Severity Index (PDSI) from the BR-DWGD and TerraClimate gridded datasets [72] using the `brclimr` R package [73]. Monthly mean UV-B irradiance was extracted from the glUV database [74] and photoperiod was calculated from geographic coordinates using the `LightLogR` R package [75]. All climate variables were aggregated to monthly means spanning the same period as the field sampling.

To characterize the temporal dynamics of each phenotypic module, **DBP** computed Euclidean distance matrices from the monthly mean values (standardized across variables) and defined trajectories in the ordination space [76, 77] in the `ecotraj` package. Three trajectory metrics were extracted: (i) trajectory length (total Euclidean distance traversed across 12 months) that measures the amount of phenotypic change, (ii) mean speed (length / 11 steps) that measures the rate of change per month, and (iii) mean angle between consecutive displacement vectors (higher angles indicate more erratic trajectories), following the framework of Phenotypic Trajectory Analysis [71]. Because modules differ in the number of variables (3–5) and monthly sample sizes (n = 1–7 for histochemistry and morphometry), we performed two sensitivity analyses: (1) recomputing trajectory metrics on PC1 × PC2 scores only to assess dimensionality effects, and (2) excluding months with n ≤ 2 individuals to assess the influence of single-specimen centroids. In both cases, we compared speed rankings via Spearman correlation.

To quantify the uncertainty of the trajectory metrics and to test whether modules differ, **DBP** implemented a paired non-parametric bootstrap [78]. Within each replicate, individuals were resampled with replacement within each month (preserving the observed monthly sample sizes), monthly centroids were recomputed, and the trajectory metrics — mean speed, path length, and directionality (the ratio of net displacement to total path length, ranging from 0 to 1) — were re-estimated for all four modules on the same resampled months, so that between-module differences are evaluated on paired replicates. For volumetric density, the complete compositional pipeline (amalgamation, count-zero-multiplicative imputation, and centred-log-ratio transformation) was repeated inside each replicate, so that its intervals are computed on the same scale as the observed metrics. We obtained percentile 95% confidence intervals (2.5th–97.5th percentiles) for each metric, and tested pairwise differences between modules with a two-sided bootstrap test (P = 2 × the smaller of the proportions of replicates in which the difference was ≤ 0 or ≥ 0). Analyses used the ecotraj [76], compositions, and zCompositions [66] packages, with a fixed random seed for reproducibility.

**DBP** quantified concordance among phenotypic modules and climate using the PROcrustean randomization TEST (PROTEST, [79, 80]). For each of the 10 pairwise comparisons among the five datasets (somatic indices, histochemistry, cell morphometry, tissue volumetric density, and climate), we computed monthly centroids (n = 12 months) and applied symmetric Procrustes superimposition on z-standardized data matrices, yielding the m² statistic (sum of squared residuals; lower values indicate stronger concordance, without assuming directionality).

Because monthly time series exhibit strong temporal autocorrelation, we replaced the standard free-row permutation with a cyclic-shift permutation scheme that preserves the autocorrelation structure under the null hypothesis [78, 81, 82]. Under this scheme, the response matrix **Y** is cyclically shifted across its full period (n = 12), and mirror reflection doubles the permutation space from n − 1 = 11 to 2(n − 1) = 22 unique permutations. The minimum attainable *P*-value is therefore 1/23 ≈ 0.043. Because Holm correction across 10 simultaneous tests would require *P* < 0.005—an impossibility under this design—we report raw *P*-values, FDR-corrected *P*-values [83], and use the m² ranking as the primary inferential summary statistic. A detailed justification and Monte Carlo validation of the cyclic-shift approach to PROTEST will be published elsewhere. A Monte Carlo simulation [78] confirmed adequate Type I error control under the cyclic-shift scheme (Additional file 2).

To test whether climate leads biological response by *k* months, **DBP** computed lagged PROTEST for temporal lags *k* = 0, 1, 2, and 3 months for each directed edge in the DAG (Figure 1). This approach also tests whether the speed of response differs across levels of biological organization. At each lag *k*, the first *n* − *k* rows of the driver matrix **X** are matched to the last *n* − *k* rows of the response matrix **Y**, and m² is computed via symmetric Procrustes rotation. A decrease in the m² statistic at lag *k* > 0 relative to lag 0 provides evidence that the driver leads the response by approximately *k* months. For each edge, both the hypothesized and reverse directions were tested, which allowed us to assess whether the relationship is directional or bidirectional. The optimal lag for each edge was identified as the lag minimizing m² in the hypothesized direction. This approach uses the full multivariate data for each module and is robust (Additional file 2) to the short time series (n = 12).

To address whether improvement in m² at higher lags could be an artifact of reduced sample size (*n* − *k* rows rather than *n*), **DBP** computed the Δm² = m²(lag 0) − m²(best lag) and compared it to a null distribution generated by applying the same cyclic-shift + mirror permutations at both lag 0 and the best lag (Additional file 2). If the observed Δm² exceeded the 95th percentile of the null distribution, it can be concluded that the improvement was genuine. Analysis was conducted in the R package `vegan` [84].

To visualize when each module undergoes its greatest rate of change, **DBP** computed the Euclidean displacement between consecutive monthly centroids in PC1 × PC2 space. This yields 11 step sizes (April → May, …, February → March) per module. These speed profiles were normalized within each module (0–1) to allow comparison of the sequence of peak change. If the hypothesized response hierarchy holds, peak speed should occur earliest in histochemistry and latest in somatic indices.

As complementary univariate evidence, **DBP** fitted simple linear models of climate PC1 on each biological PC1, comparing the contemporary model (lag 0; n = 12) with the lagged model (climate at *t* − 1 predicting biology at *t*; n = 11). An increase in R² at lag 1 supports the interpretation that climate leads the biological response by approximately one month. Analysis was conducted in the R package `easystats` [85]. All analyzes were conducted in R v. 4.5.2 [86].

## Results

Climate variables showed a clear seasonal pattern along the first two principal components, with PC1 (72.3% of variance) representing the warm/wet vs. cold/dry gradient and PC2 (21.0%) capturing variation in relative humidity relative to maximum temperature (Fig. 2). The dry season (June–August) clustered on the negative end of PC1, characterized by low temperature, rainfall, UV-B, and photoperiod, but high drought severity (PDSI). The wet season (November–March) occupied the positive end, with the reverse loadings. April and September represented transitional periods. The PCA of each phenotypic module captured substantial proportions of total variance in two axes. All modules exhibited clear seasonal trajectories in ordination space, with months arranged in a broadly cyclical pattern from dry to wet season (Fig. 2).

**Figure 2.**
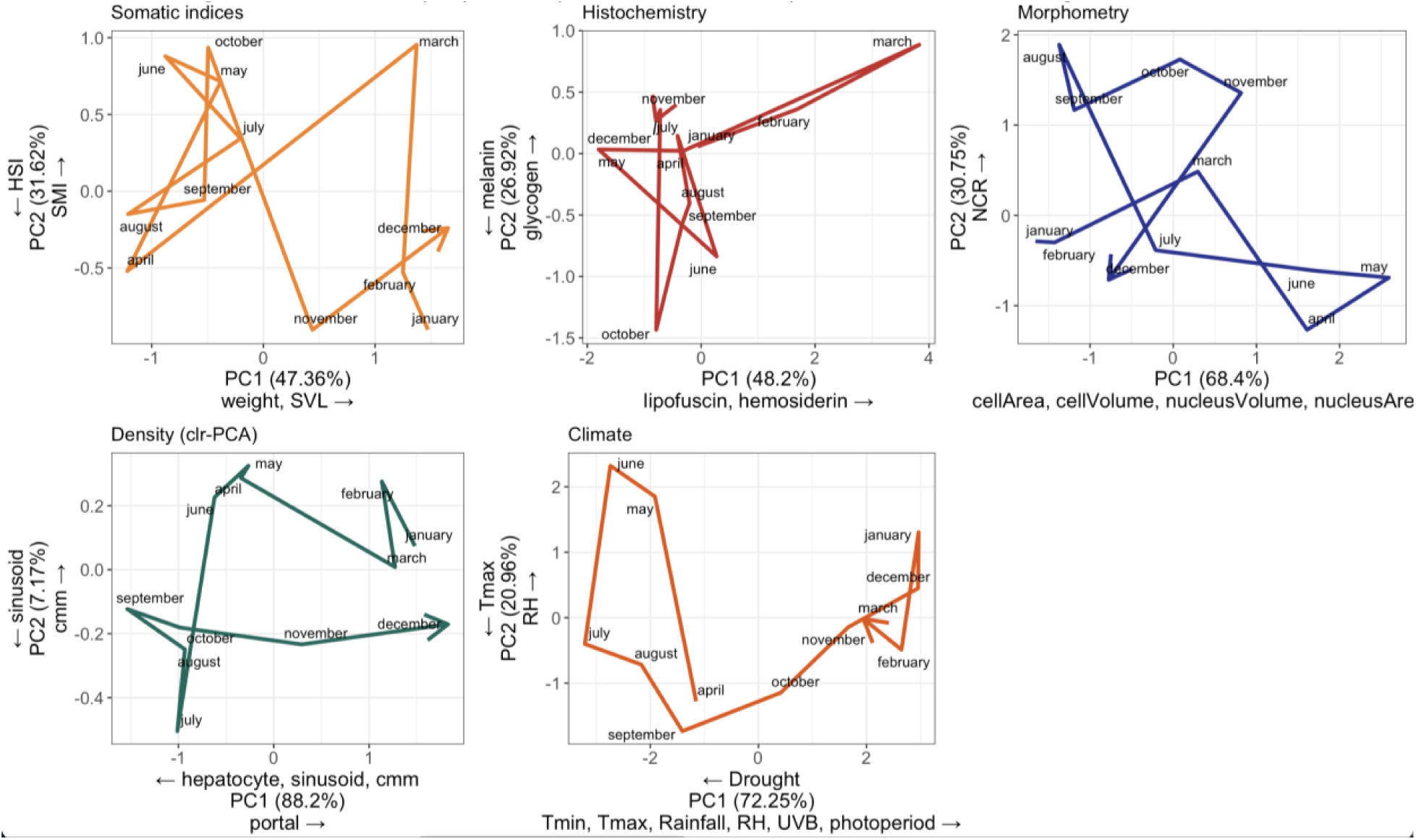
Seasonal phenotypic trajectories for each module and climate in PC1 × PC2 ordination space. Monthly centroids are connected by arrows indicating the direction of temporal change (April 2016 → March 2017). The variables with the strongest loadings (|r| > 0.5) in each direction.

Somatic indices, histochemistry, and morphometry formed a fast-responding group (mean speeds 1.98–2.39), while volumetric density was distinctly slower (mean speed 0.76; Table 1; Additional file 1: Fig. S4). Bootstrap 95% confidence intervals [78] confirmed this structure (Additional file 1: Table S3). In a paired bootstrap that resampled individuals within months so that all four modules shared the same 12-month structure in each replicate, volumetric density differed significantly from each of the three fast modules in both speed (differences of 1.22–1.34 ED per month, all P < 0.001) and path length (differences of 13.4–14.8 ED, all P < 0.001), whereas the three fast modules did not differ from one another in either metric (all pairwise P > 0.55). Directionality did not differ between any pair of modules (all P ≥ 0.30), indicating similarly tortuous trajectories throughout (Additional file 1: Table S4). The two-tier speed structure is therefore robust, whereas the fine-grained ordering within the fast group is not statistically resolved (Additional file 1: Fig. S3).

**Table 1.**
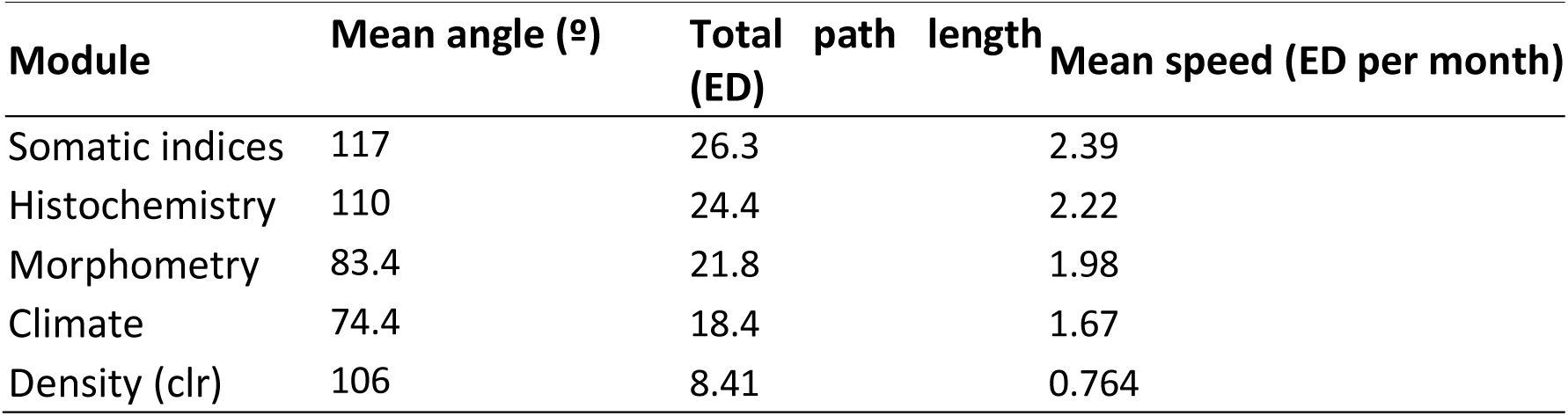
Trajectory metrics for each phenotypic module and climate. ED = units of Euclidean Distance.

| Module | Mean angle (°) | Total path length (ED) | Mean speed (ED per month) |
| --- | --- | --- | --- |
| Somatic indices | 117 | 26.3 | 2.39 |
| Histochemistry | 110 | 24.4 | 2.22 |
| Morphometry | 83.4 | 21.8 | 1.98 |
| Climate | 74.4 | 18.4 | 1.67 |
| Density (clr) | 106 | 8.41 | 0.764 |

A within-month coefficient of variation (CV) sensitivity analysis ruled out the possibility that differential measurement noise drives the speed ranking (Additional file 1: Fig. S2). Somatic indices, the fastest module, had the lowest within-month CV (median = 0.13) comparable with morphometry (median = 0.121) – indicating high measurement precision –while histochemistry (median = 0.506) and density (median = 0.498) had the highest, despite being the slowest and second-fastest modules, respectively. This inverse relationship between speed and CV confirms that the observed speed differences reflect genuine biological response dynamics rather than measurement artifacts. However, two sensitivity analyses revealed that the ranking within the fast group is not fully stable (Additional file 1: Fig. S3). First, recomputing trajectory metrics on PC1 × PC2 scores alone yielded Spearman ρ = 0.2 with the full-variable ranking (dimensionality sensitivity). Second, excluding months with n ≤ 2 individuals for histochemistry and morphometry (7 months retained) also produced ρ = 0.2 (low-n sensitivity): morphometry became fastest (2.90), followed by histochemistry (2.26) and somatic indices (2.09), while density remained slowest (0.76). In both cases, the qualitative separation between the fast-responding group and the slow density tier was preserved, but the within-group ordering is sensitive to methodological choices [87]. We therefore interpret the fast/slow dichotomy as robust, while the fine-grained ranking among somatic indices, histochemistry, and morphometry should be treated with caution.

Mean trajectory angles further differentiated the modules (Table 1). Climate displayed the lowest mean angle (74.4°), consistent with a smooth, predictable seasonal cycle. Among the biological modules, morphometry was the most directional (83.4°), suggesting that cellular changes track a gradual, unidirectional seasonal signal. In contrast, somatic indices (117°) and histochemistry (110°) exhibited obtuse mean angles, indicating frequent reversals in the direction of phenotypic change between consecutive months — consistent with their sensitivity to short-term fluctuations in food availability and hydration. Volumetric density combined a high mean angle (106°) with the lowest speed (0.76), indicating minimal, but erratic change in tissue composition.

The strongest contemporaneous (Fig. 3) concordance was between Density and Climate (m² = 0.378, raw *P* = 0.087, FDR *P* = 0.290) and between Somatic indices and Climate (m² = 0.476, raw *P* = 0.043, FDR *P* = 0.290). Histochemistry showed moderate concordance with Density (m² = 0.704) and Climate (m² = 0.791). The weakest climate concordance was with Morphometry (m² = 0.771), and the weakest concordances overall were among the biological modules, notably between somatic indices and morphometry (m² = 0.815). Unexpectedly, histochemistry and morphometry were also among the least concordant pairs (m² = 0.806). These results indicate that the four phenotypic modules are largely temporally independent. Given the *P*-value floor of 0.043 imposed by the cyclic-shift permutation space, none of the 10 pairwise comparisons survived FDR correction at α = 0.05. Therefore, the m² ranking (Additional file 1: Table S1) serves as the primary summary of concordance strength (Fig. 3).

**Figure 3:**
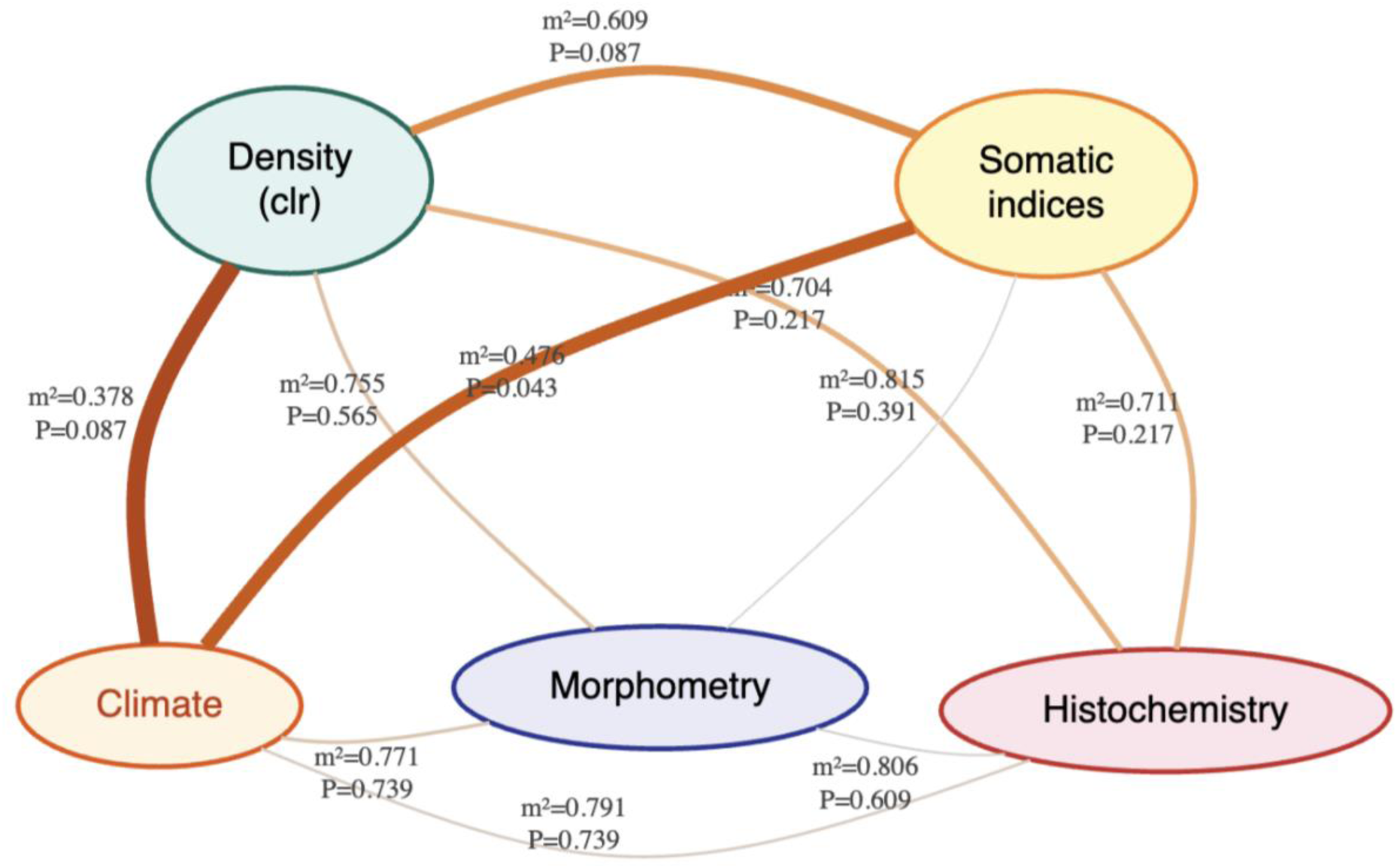
Contemporaneous PROTEST concordance network among phenotypic modules and climate. Edge color and thickness encode the Procrustean m² statistic, with thicker, darker edges indicating stronger concordance (lower m²). Edge labels report m² and the raw cyclic-shift p-value.

There was a clear temporal hierarchy in climate–biology concordance (Fig. 4, Table 2, S1). Somatic indices and volumetric density achieved their best fit with climate at lag 0 (m² = 0.476 and 0.378, respectively), indicating contemporaneous or near-instantaneous tracking. In contrast, histochemistry showed a 3-month cumulative delay (m² improving from 0.791 at lag 0 to 0.379 at lag 3, *P* = 0.043), and morphometry showed its best fit at lag 2 (m² = 0.727), though with weaker overall concordance (Table 2, S1).

**Figure 4:**
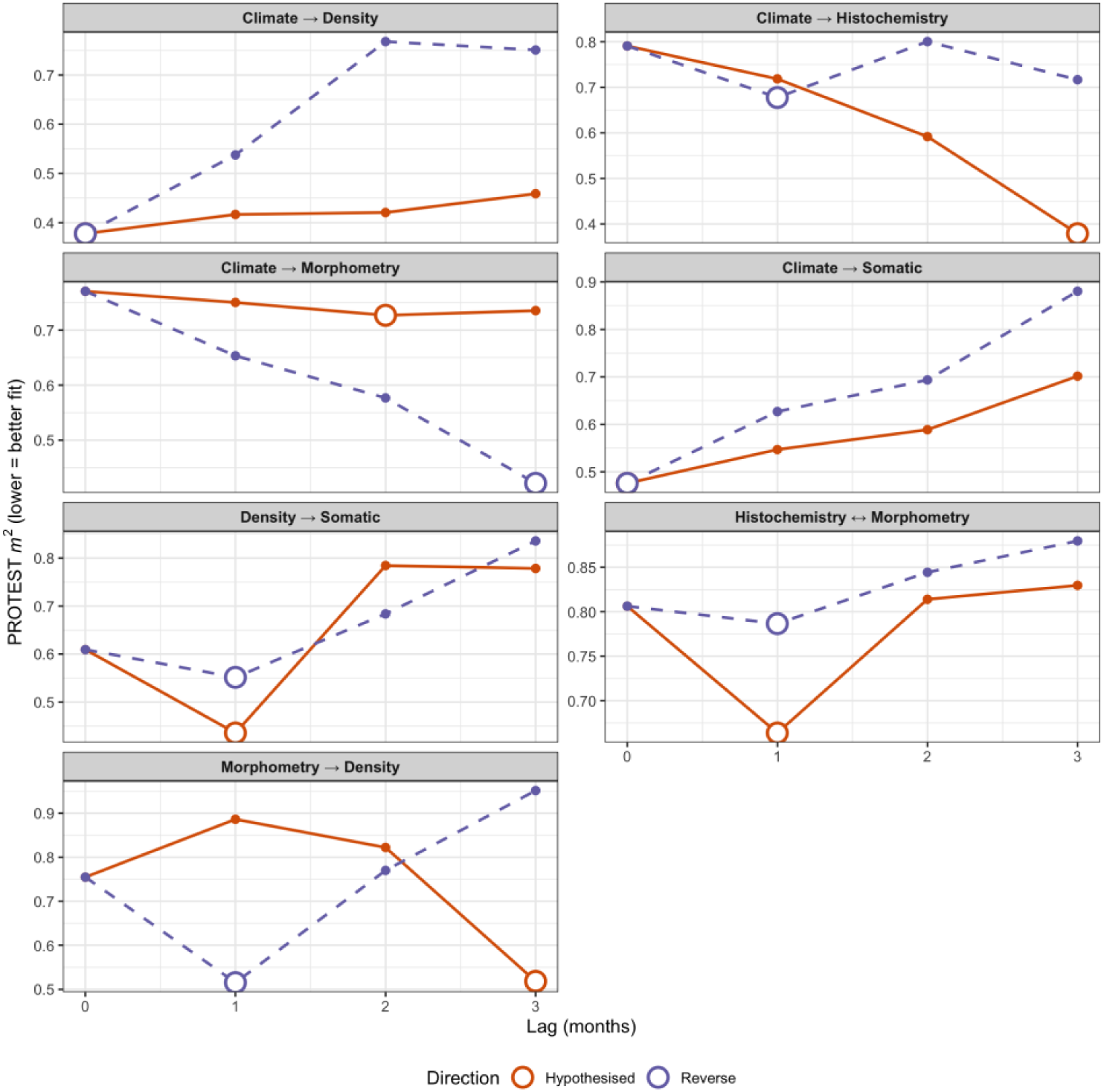
Lagged PROTEST m² across temporal lags 0–3 months, for each DAG edge in Fig. 1. Lower m² indicates stronger concordance. Solid lines = hypothesized direction; dashed = reverse. The large open circle in each panel-direction marks the lag with the minimum m² (the recovered optimal lag). Cyclic-shift permutation imposes a floor of *P* ≥ 0.043 with n = 12, so formal significance markers are omitted; m² minima are interpreted as point estimates of the lead-lag structure.

**Table 2.**
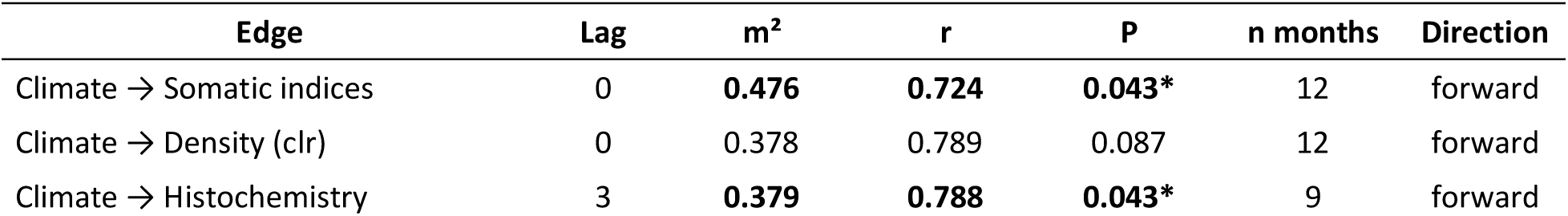
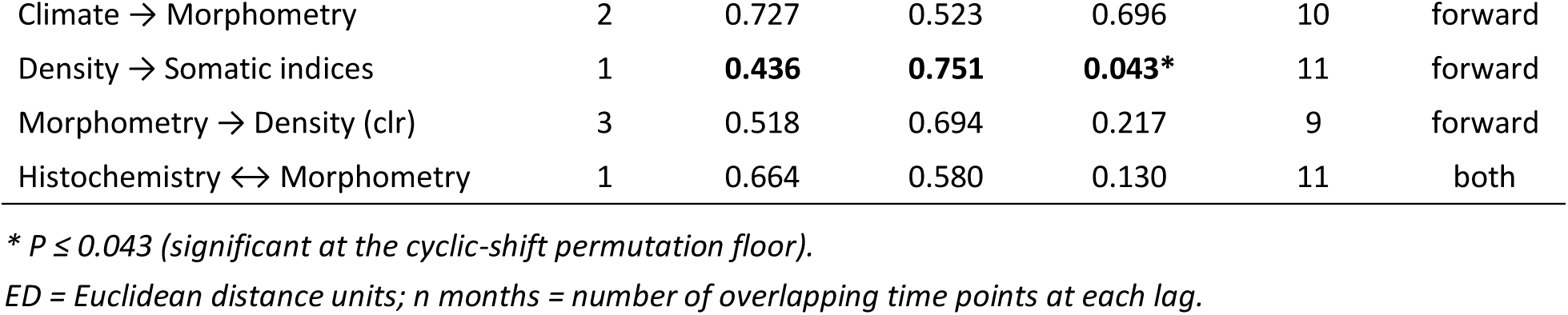
Summary of lagged cyclic-shift PROTEST results. For each edge in the hypothesized cascade (Fig. 1), the table shows the lag (in months) that minimized m² (maximized Procrustean correlation r). Only forward-direction lags are shown (climate or upstream module leading); the bidirectional edge Histochemistry ↔ Morphometry was tested in both directions, and the better fit is reported. Significance was assessed by cyclic-shift permutation (23 permutations; minimum attainable P = 1/23 ≈ 0.043). Full results for all lags and directions are in Additional file 1: Table S1.

| Edge | Lag | $m^2$ | $r$ | P | n months | Direction |
| --- | --- | --- | --- | --- | --- | --- |
| Climate → Somatic indices | 0 | **0.476** | **0.724** | **0.043\*** | 12 | forward |
| Climate → Density (clr) | 0 | 0.378 | 0.789 | 0.087 | 12 | forward |
| Climate → Histochemistry | 3 | **0.379** | **0.788** | **0.043\*** | 9 | forward |
| Climate → Morphometry | 2 | 0.727 | 0.523 | 0.696 | 10 | forward |
| Density → Somatic indices | 1 | **0.436** | **0.751** | **0.043\*** | 11 | forward |
| Morphometry → Density (clr) | 3 | 0.518 | 0.694 | 0.217 | 9 | forward |
| Histochemistry ↔ Morphometry | 1 | 0.664 | 0.580 | 0.130 | 11 | both |

\* $P \leq 0.043$ (significant at the cyclic-shift permutation floor).
ED = Euclidean distance units; $n$ months = number of overlapping time points at each lag.

Among biological inter-module links, histochemistry led morphometry by 1 month (m² = 0.664 at lag 1), and density led somatic indices by 1 month (m² = 0.436, *P* = 0.043). The Δm² permutation test confirmed that the improvement for Climate → Histochemistry at lag 3 exceeded the null expectation from sample-size reduction alone, supporting a genuine 3-month delay rather than an artifact of using n − 3 = 9 time points (Table 2).

The temporal sequence of peak displacement was consistent with the lagged PROTEST results. Climate showed its maximum rate of change in the transition from the dry to the wet season (Fig. 5). Among biological modules, histochemistry peaked earliest (consistent with its role as a cumulative integrator requiring metabolite build-up over 3 months), followed by somatic indices, cell morphometry, and finally volumetric density (Fig. 5). The heatmap of normalized speed (Additional file 1: Fig. S4) visualizes this temporal response hierarchy, with each module’s peak displacement offset in time relative to the shared climatic driver.

**Figure 5:**
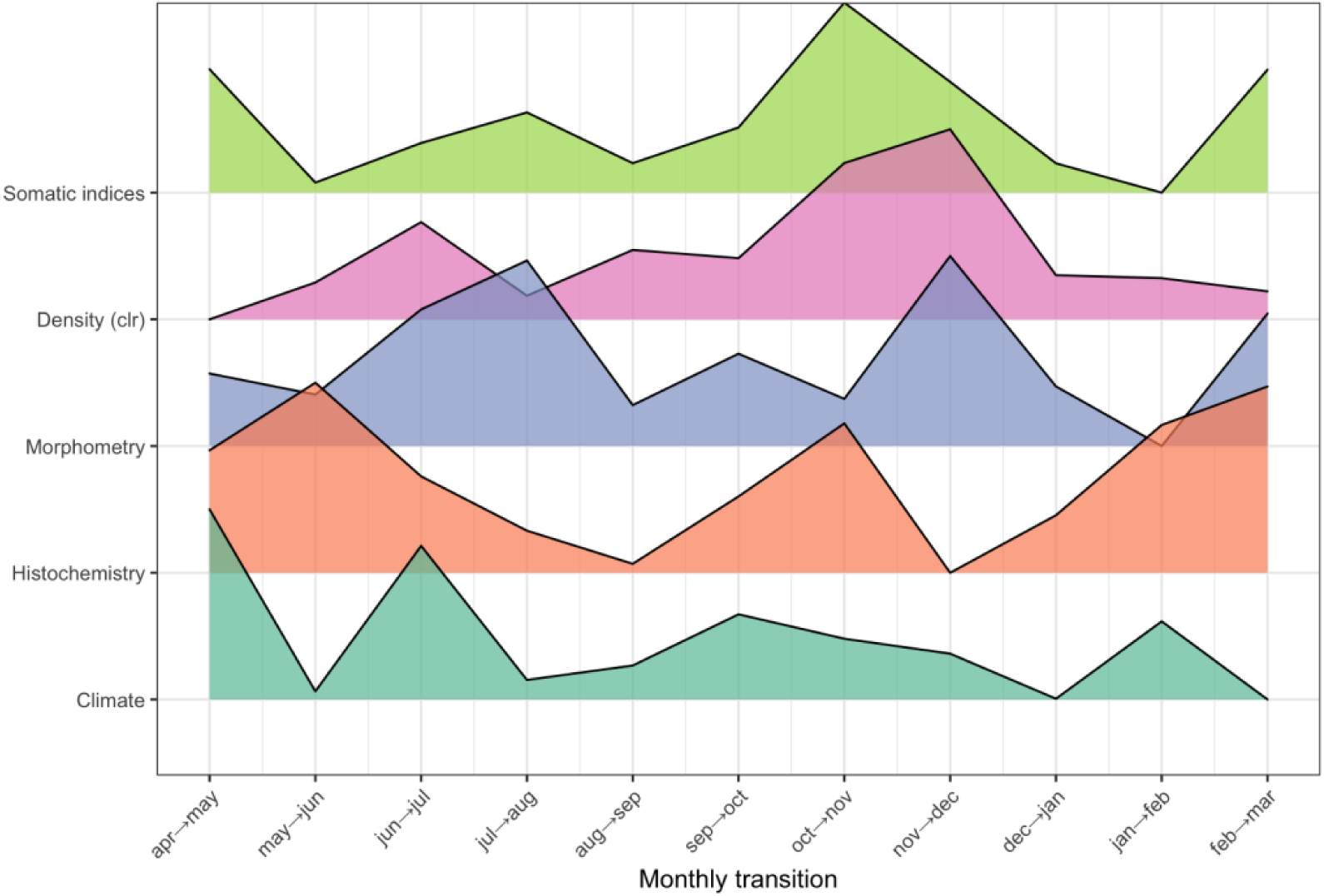
Speed profiles for each phenotypic module across the year. Peaks indicate when each module undergoes its greatest rate of change.

Climate PC1 at *t* − 1 explained more variance in histochemistry PC1 at *t* (lagged R² > contemporary R²), consistent with a one-month lead effect. For somatic indices and density, the contemporary model (lag 0) yielded higher R², corroborating the contemporaneous tracking detected by lagged PROTEST. Histochemistry and cell morphometry showed weaker relationships with climate PC1 (R² < 0.25), consistent with the PROTEST results indicating that their multivariate response is not well captured by a single principal component. With only *n* = 11–12 observations, individual regressions had low statistical power; we interpret the pattern of R² differences across modules rather than individual significance tests (Fig. 6).

**Figure 6:**
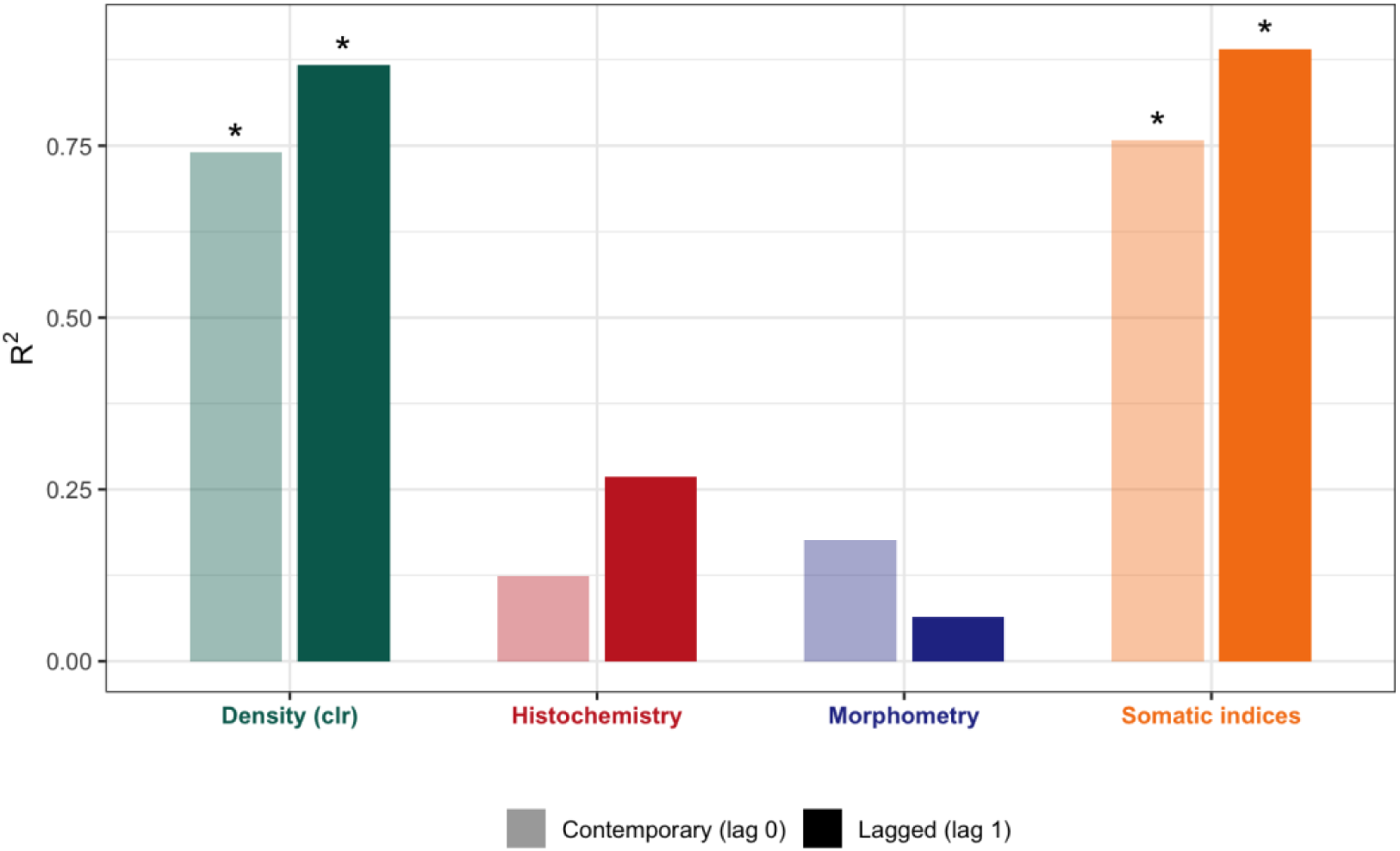
Comparison of R² between contemporary (lag 0) and lagged (lag 1) simple regressions of climate PC1 on each biological PC1. Higher R² at lag 1 indicates that climate leads the biological response by approximately one month.

The recovered lag structure (Fig. 7A) contradicts the bottom-up prediction of Bayne et al. [38]. Rather than intracellular processes (histochemistry) responding fastest, we found a descending (top-down) temporal response hierarchy: somatic indices and volumetric density tracked climate contemporaneously (lag 0), while histochemistry lagged by 3 months. Trajectory speed analysis corroborated this pattern: somatic indices (speed = 2.39), histochemistry (2.22), and morphometry (1.98) formed a fast-responding group with overlapping 95% bootstrap confidence intervals, whereas density (speed = 0.76) constituted a separate slow tier (Fig. 7B). Importantly, trajectory speed (amount of change per unit time) and PROTEST lag (temporal delay before concordance peaks) represent complementary, orthogonal axes of the temporal response (Fig. 7B): a module can change rapidly, yet with a delayed onset (as observed for histochemistry), or slowly but contemporaneously (as observed for volumetric density).

**Figure 7.**
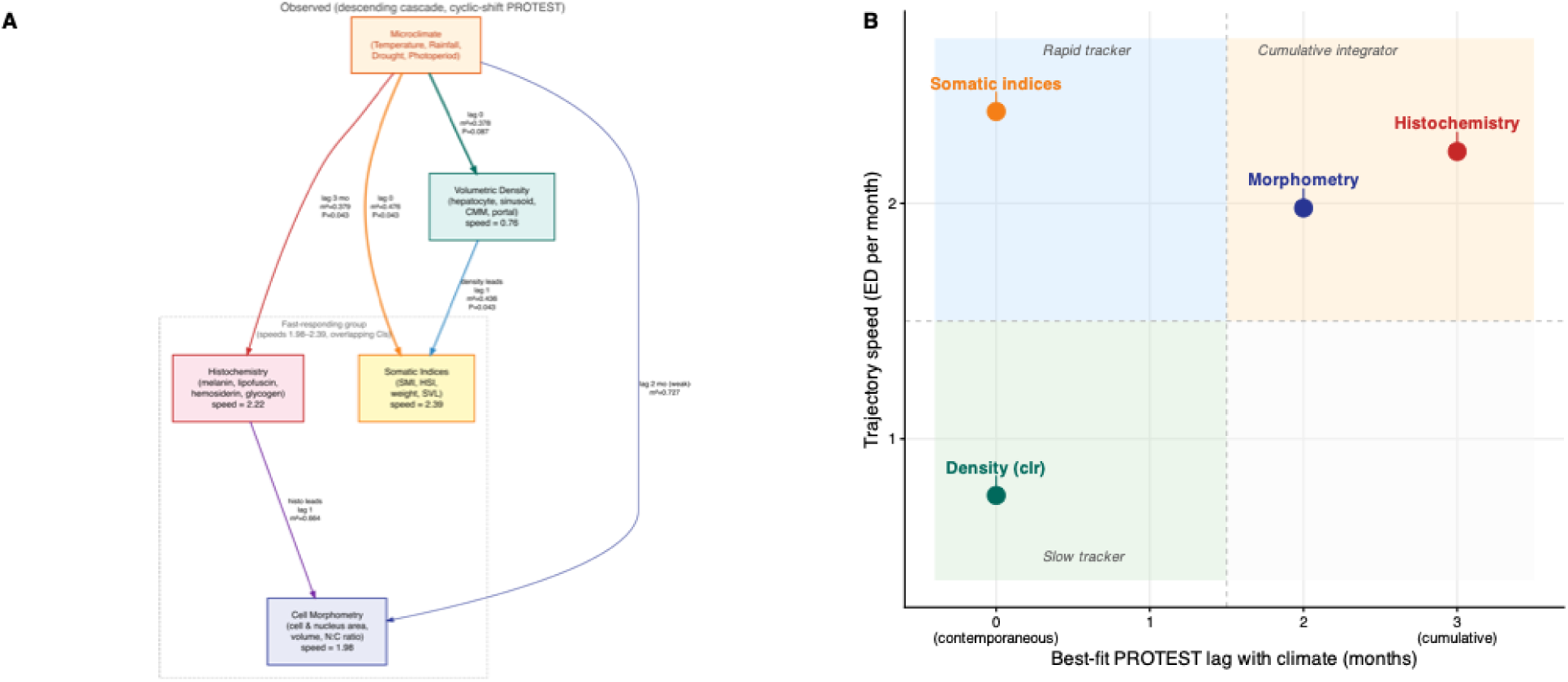
Observed temporal response hierarchy of phenotypic modules to climate. **(A)** Directed acyclic graph summarizing the cyclic-shift PROTEST results. Microclimate sits at the top; arrows point to each biological module at the optimal lag (m² minimum). Edge labels report the best-fit lag, m², and raw permutation *P*; edge thickness is proportional to concordance strength (1 − m²). The dashed bracket groups modules whose bootstrap confidence intervals for trajectory speed overlap (fast-responding group), while volumetric density forms a separate slow tier. **(B)** Response-mode landscape. Each module is positioned by its best-fit PROTEST lag (Table 2) with climate (*x*-axis) and its mean trajectory speed in PC1–PC2 space (*y*-axis). The dashed lines partition the space into three response modes: *rapid trackers* respond contemporaneously and fast (somatic indices), *cumulative integrators* accumulate a multi-month climatic signal, yet still change rapidly (histochemistry, morphometry), and *slow trackers* respond contemporaneously, but with low displacement per month (volumetric density).

## Discussion

Our results reveal a two-tier temporal structure in the seasonal phenotypic response that contrasts with the bottom-up cascade predicted by Bayne et al. [38]. Rather than intracellular processes responding first, somatic indices and volumetric density tracked climate contemporaneously (PROTEST lag 0), whereas histochemistry and morphometry required a three-month accumulation window (lag 3) to reach peak concordance with climate. Trajectory speed further partitioned the modules: somatic indices, histochemistry, and morphometry formed a fast-responding group, while density constituted a distinct slow tier. These two axes — temporal lag and displacement rate — are complementary: a module can change rapidly yet with delayed onset (histochemistry), or contemporaneously yet with minimal displacement (density). This complementarity resonates with the recent finding that, across ectothermic species, high capacity for physiological plasticity is associated with a slow rate of phenotypic change [88], suggesting that the rate–capacity trade-off documented at the interspecific level also manifests among phenotypic modules within a single organism. More broadly, thermal responses seem to differ systematically across levels of biological organization in ectotherms [89], with higher organizational levels increasing more rapidly than lower ones, consistent with the temporal response hierarchy we report here. We propose that this structure reflects a temporal response hierarchy in which seasonal climate enters the organism through whole-body energy balance and hormonal signaling, with downstream effects on intracellular biochemistry emerging only after sustained metabolic shifts.

This reversal reflects how the driver enters the organism. Bayne et al.’s (1985) bottom-up cascade was originally formulated for toxicant exposure in marine invertebrates, where chemicals enter cells directly through membrane transport, perturbing intracellular processes before effects propagate upward to tissues and the whole organism. Seasonal climate, by contrast, acts primarily through whole-organism physiological pathways: temperature affects metabolic rate globally [7], while precipitation determines prey availability [55] and thus energy intake. These pathways first alter energy balance and hormonal status (organismal level), with downstream effects on cellular biochemistry emerging only after sustained shifts in metabolic demand.

Somatic indices responded contemporaneously with climate and were the fastest-responding module, consistent with whole-organism indices being the first to register environmental change at monthly resolution. The high trajectory angle further supports this: body mass and condition of *D. minutus* respond not only to the overall seasonal trend [14, 16], but also to short-term pulses in rainfall and food availability [7, 19], producing frequent directional reversals in the ordination space. Even muscular metabolic biomarkers in seasonally active anurans respond in a species-specific manner to seasonal fluctuations [35], underscoring the tight coupling between organismal energy status and environmental conditions. This contemporaneous response reflects the direct effects of temperature on metabolic rate and food availability on energy intake, both of which translate rapidly into changes in body mass and liver mass [39, 51]. The liver’s role as the primary glycogen and lipid storage organ in frogs means that hepatosomatic index fluctuations are among the most immediate physiological consequences of seasonal climate variation [22, 50]. In contrast, morphometry was the most directional module, suggesting that cellular changes track a gradual, seasonal signal with fewer reversals. This finding resonates with Wu et al. [90], who found no evidence that lower-level biological responses are induced faster than higher-level ones in the context of pollution exposure. Our study extends this perspective to natural seasonal variation, showing that when the environmental driver acts through whole-organism physiology rather than direct intracellular perturbation, the temporal hierarchy is reversed.

Volumetric density responded contemporaneously with climate, yet its trajectory speed was threefold lower than that of any other module. This combination of strong concordance but low displacement defines density as a *slow tracker* (Fig. 7B): tissue composition faithfully mirrors the direction of climatic variation, but the magnitude of month-to-month change is small. This pattern suggests that hepatic tissue architecture functions as a structurally conservative compartment, buffering short-term seasonal fluctuations while maintaining overall organ integrity. The relative proportions of hepatocytes, sinusoids, and melanomacrophage centers reflect a structural equilibrium that shifts only incrementally because tissue remodeling requires coordinated cell proliferation and extracellular matrix turnover [30, 31], which may take days to weeks to happen. Seasonal hepatic adjustments of this kind appear conserved across ectothermic vertebrates: hepatocyte and nuclear volumes change in brown trout (*Salmo trutta*) in a stage-dependent manner across the reproductive cycle, with cell turnover markers (PCNA, caspase-3) confirming that tissue remodeling is gradual rather than abrupt [91]. Hepatocytes become larger and more vacuolated in piranhas (*Pygocentrus nattereri*) during the rainy season, paralleling the increased metabolic activity signaled by elevated ALT and triglycerides [92]; and in overwintering lizards (*Uta stansburiana*), hepatic glycogen, instead of lipids, limits winter survival, underscoring the liver’s role as a seasonally regulated energy buffer [93]. In tegu lizards (*Tupinambis merianae*), seasonal metabolic depression involves a 50% depletion of liver glycogen alongside differential redistribution to heart and brain [94], illustrating that hepatic reserves can be mobilized at organ-specific rates, a parallel to the module-specific timescales we document here. The circannual rhythm of liver tissue composition documented in hibernating frogs [32] thus appears to operate in tropical non-hibernating populations as well, albeit with smaller amplitude and no discrete hibernation-linked discontinuity. Comparable seasonal hepatic adjustments have also been described in another year-round active Neotropical hylid, *Boana prasina*: winter individuals had larger livers, but lower hepatic citrate synthase activity, indicating that organ-mass compensation operates alongside reduced specific aerobic capacity [95]. Notably, density led somatic indices by one month, suggesting that subtle shifts in tissue architecture precede the more conspicuous changes in organ mass and body condition.

The histochemical module showed a progressive improvement in concordance from lag 0 to lag 3, suggesting a cumulative rather than immediate response to climate. This is consistent with the biology of liver pigments and metabolites. Glycogen reserves accumulate gradually during favorable periods and are depleted during energetically demanding periods, such as reproduction [23, 28, 50], while melanin and lipofuscin deposition in melanomacrophage centers reflects cumulative oxidative damage and immune activity over weeks to months [32, 47]. The circannual rhythm of hepatic melanin [40] supports this multi-month integration, and hepatic pigments also respond to thermal stress [54] and land-use change [48] in frogs, reinforcing their role as integrators of environmental conditions across timescales. Similarly, the temporal patterns of lipid labeling in liver tissue [41] demonstrate that lipid metabolism in the frog liver follows seasonal patterns with characteristic delays relative to environmental cues. Taken together, our results indicate that hepatic hemosiderin and lipofuscin may function as seasonally integrated biomarkers in tropical anurans: rather than responding exclusively to acute variation, these pigments accumulate progressively across seasonal windows, integrating fluctuations in metabolic activity, erythrocyte turnover, oxidative stress, and immune function, converging with the recognized role of melanomacrophage pigments as long-term physiological archives in ectothermic vertebrates [47]. Under this perspective, hemosiderin and lipofuscin may provide not only indicators of environmental stress [47, 48], but also markers of seasonal physiological allocation and energetic history [25, 32]. These contrasting temporal signatures — near-instantaneous coupling for somatic indices vs. progressively delayed concordance for histochemical traits — indicate that climate forcing propagates through the hepatic system at module-specific timescales. Notably, this cumulative signal emerged clearly in the multivariate analysis, but only weakly when histochemistry was reduced to its first principal component (∼48% of variance), suggesting that the climate response is distributed across multiple pigments and metabolites rather than concentrated along a single dominant axis, which is consistent with different histochemical markers tracking different facets of environmental forcing (energy balance, oxidative stress, immune activity).

Histochemistry consistently preceded cell morphometry, while the reverse direction worsened. This temporal precedence is consistent with the expectation that intracellular biochemical changes (pigment accumulation, glycogen deposition) should induce subsequent changes in cell size and shape [9, 25, 49]. Frog hepatocytes become hypertrophic with large amounts of storage material during hibernation preparation [25]. However, cell swelling itself promotes glycogen synthesis via activation of glycogen synthase [96], creating a positive feedback loop. The weak contemporaneous concordance between histochemistry and morphometry suggests that intracellular metabolite accumulation and cellular hypertrophy may operate at finer temporal scales than those captured by monthly sampling intervals. The improvement in concordance with increasing lag is consistent with metabolite accumulation initiating the process and cell volume changes following as a secondary consequence.

The two strongest pairwise PROTEST associations both involved climate: climate–density and climate–somatic indices, indicating that these modules most closely track climate. In contrast, all purely biological pairs exhibited weak concordance, with only density-somatic indices showing moderate coupling. This pattern is consistent with Bayne et al.’s (1985) prediction that adjacent levels of biological organization should be more tightly linked than distant ones. The predominant temporal independence among biological modules we found is consistent with phenotypic modularity theory but goes further: recent work shows that phenotypic integration does not constrain the magnitude of plastic responses; instead, it limits their variation, producing correlated plasticities among tightly integrated traits [97, 98]. In the liver, the observed independence among modules implies that each is governed by distinct proximate physiological mechanisms with different intrinsic timescales: melanomacrophage dynamics respond to seasonal changes in oxidative stress and melanosynthetic activity with a time course set by cell proliferation and apoptosis cycles [32], whereas somatic indices, such as HSI and body mass track the more immediate metabolic balance between energy intake and expenditure [28]. The weak concordance between histochemistry and morphometry – despite their known mechanistic coupling via the glycogen-cell volume feedback loop [96] – might suggest that intracellular hypertrophy may operate at finer temporal scales than those captured by the monthly sampling intervals (see [23]). This modular structure means that differential plasticity among modules can reorganize trait covariation patterns across environments [97], a process that may be especially relevant in strongly seasonal systems, in which the direction and intensity of environmental forcing change cyclically. The density–somatic coupling (the strongest biological pair) is expected on mechanistic grounds, since hepatocyte vacuolization — a component of tissue volumetric density — directly reflects lipid and glycogen storage, which in turn determines organ mass and condition [21, 28]. The fact that this pair does not reach formal significance underscores the low power inherent to 12-point time series, rather than absence of a biological relationship.

Our study has several limitations. First, we sampled only adult males; females experience different seasonal pressures (vitellogenesis, egg production, and associated hepatic lipid mobilization) that profoundly alter liver physiology [18]. The temporal response hierarchy we document may differ in females, where reproductive demands could accelerate or delay specific modules relative to the pattern we observed in males. Second, the study is based on a single population sampled over one annual cycle, which limits the generalizability of the specific lag values and phase timings. The bootstrap confidence intervals for trajectory metrics partially address sampling uncertainty, but inter-annual variation remains untested. Third, monthly sample sizes for the histochemistry and morphometry modules (n = 1–5 per month, 40 total) are small, and months with n = 1–2 yield centroids of limited reliability. We mitigated this through bootstrap resampling [78], but future studies should aim for more balanced sampling designs. Multi-year sampling would allow testing whether the two-tier speed structure is a robust, repeatable feature or whether it varies with climatic conditions. Additionally, our correlative design cannot definitively establish that climate causes the observed phenotypic changes; experimental manipulation would be needed to confirm the causal directions implied by the lagged PROTEST results.

A further consideration is that we sampled adult males throughout their reproductive season, so male reproductive investment could contribute to the hepatic variation we attribute to climate. Advertisement calling is energetically expensive in anurans [99, 100], and in Neotropical assemblages calling activity is concentrated in the warm, rainy months and covaries with the same temperature and rainfall variables that drive energy intake [101, 102]. Spermatogenesis in *D. minutus* and other Neotropical anurans likewise follows a seasonal germinative cycle [17, 18, 103]. Because reproductive effort therefore covaries with climate, our observational design cannot fully disentangle reproductive from climatic drivers: the temporal response hierarchy we describe is a pattern relative to measured climatic variables, and unmeasured reproductive investment could contribute to or modify it. Incorporating reproductive metrics — such as calling activity and a gonadosomatic index — into future multi-year sampling would help separate these coupled drivers.

Extending this framework to other species and organ systems would test whether the temporal response hierarchy is a general feature of seasonal plasticity or specific to hepatic physiology. Comparative studies across species with contrasting life histories [28, 35] would be particularly informative, especially considering the growing recognition that the rate of phenotypic plasticity — not just its magnitude — is a key determinant of organismal resilience under environmental change [2, 3, 105]. Quantifying half-times and displacement rates for hepatic modules across species would reveal whether the rate–capacity trade-off documented at the interspecific level [88] also structures intraorganismal plasticity more broadly. This pronounced modularity has practical implications for biomonitoring: seasonal changes in one aspect of liver physiology cannot be reliably predicted from another, and single-biomarker approaches may miss relevant phenotypic variation. Franco-Belussi et al. [48] reached a similar conclusion from the idiosyncratic species-level pigment responses to land use change. Future studies combining our temporal plasticity framework with quantitative genetic data could test whether the hepatic modules that respond fastest also harbor the greatest additive genetic variation (*V_A_*), as predicted by Noble et al. [104], which would indicate high evolutionary potential (*V_A_>>0*) in these traits. Finally, applying the same approach to urbanization gradients (e.g., [46]) could test whether chronic anthropogenic stress shifts the cascade from top-down back toward the bottom-up pattern, as predicted by Bayne et al. [38].

## Conclusions

In a wild, non-hibernating tropical frog, seasonal phenotypic plasticity of the liver follows a temporal response hierarchy rather than the bottom-up cascade predicted for toxicant exposure: whole-organism energy balance and tissue composition track seasonal climate contemporaneously, whereas intracellular histochemistry and cell morphometry lag by about three months. The direction of the hierarchy therefore depends on how the environmental signal enters the organism — through whole-body physiology for seasonal climate rather than through direct molecular disruption. The lagged, cyclic-shift PROTEST framework introduced here offers a general, simulation-validated tool for testing temporal precedence in short, autocorrelated multivariate biological time series, and can be extended to other organs, taxa, and environmental drivers.

## Declarations

### Ethics approval and consent to participate

This study was approved by the Institutional Animal Care and Use Committee (CEUA, protocol #150/2015) and complied with resolution #34/2017 of the Brazilian National Council for the Control of Animal Experimentation (CONCEA) and the AVMA Guidelines for the Euthanasia of Animals (2020). Specimens were collected under permit #18573-2 issued by the Instituto Chico Mendes de Conservação da Biodiversidade (ICMBio). This study did not involve human participants, so consent to participate is not applicable.

### Consent for publication

Not applicable.

### Availability of data and materials

Raw histological micrographs will be deposited in MorphoBank upon acceptance. All data and reproducible R code (Quarto document) used to conduct the analyzes reported in this study are publicly available on Zenodo (https://doi.org/10.5281/zenodo.20301880).

### Authors’ contributions

**Lilian Franco-Belussi**: Data curation (equal), Investigation (equal), Writing – original draft (equal), Validation (equal), Methodology (equal), Writing – original draft (contributing), Writing – review & editing (equal). **Luciana T. Moraes**: Data curation (equal), Investigation (equal), Validation (equal), Writing – review & editing (equal). **Classius De Oliveira**: Conceptualization (equal), Funding acquisition (lead), Methodology (equal), Project administration (equal), Resources (equal), Supervision (equal), Writing – review & editing (equal). **Carlos E Fernandes:** Conceptualization (equal), Funding acquisition (contributing), Methodology (equal), Project administration (equal), Resources (equal), Supervision (equal), Writing – review & editing (equal). **Diogo B Provete**: Conceptualization (lead), Formal analysis (lead), Writing – original draft (equal), Writing – review & editing (equal), Software (lead), Visualization (lead).

## Acknowledgements

Thiago Gonçalves-Souza, Kimberly Thompson, and Miquel De Cáceres provided feedback on data analysis and visualization. Karine F. Nogueira, Bruno S. L. Valverde, and Taynara Leão helped with data acquisition in the lab and field work.

## AI Use Statement

This section follows the AIdIT standard (Drobniak et al. 2026).

*Data analysis design.* Claude (Anthropic, claude-Opus-4-20250514) was consulted to evaluate the appropriateness of analytical methods, including the cyclic-shift permutation scheme for PROTEST and the compositional data analysis workflow for volumetric density. All analytical decisions were made by DBP based on the AI suggestions and independently verified against published methodological literature.

*Literature search and verification.* The AI tool was used to search for and verify relevant references, particularly for the compositional data analysis framework and the lagged PROTEST methodology. All citations were independently verified by DBP using Google Scholar and the original publications.

*Drafting and conceptual synthesis.* The AI assisted in structuring and drafting the Discussion, particularly in articulating the contrast between Bayne et al.’s (1985) bottom-up framework and our observed temporal response hierarchy. All text was reviewed, revised, and approved by all authors.

*Figure design.* The AI assisted in designing the DiagrammeR-based DAGs (Figs. 1 and 7A), the PROTEST concordance network (Fig. 3), and the speed × lag scatter plot (Fig. 7B).

*Human Oversight.* The authors maintained continuous, active oversight of all AI-generated content. Every output was critically evaluated, revised where necessary, and verified against primary sources before inclusion.

## Additional files

Additional file 1 — Supplementary Material (PDF): Figures S1–S8 and Tables S1–S4, including the dataset structure, within-month CV and low-n sensitivity analyses, bootstrap confidence intervals and paired-bootstrap tests of the trajectory metrics, the normalized-speed heatmap, representative photomicrographs of each histological module, the full lagged PROTEST results, and monthly sample sizes.

Additional file 2 — Appendix S1 (PDF): Monte Carlo simulation validating the Type I error control and lag recovery of the cyclic-shift lagged PROTEST framework.

