## Supplementary material for "Seasonal hepatic plasticity follows a temporal response hierarchy in a Neotropical frog": Online Supplementary Material 1

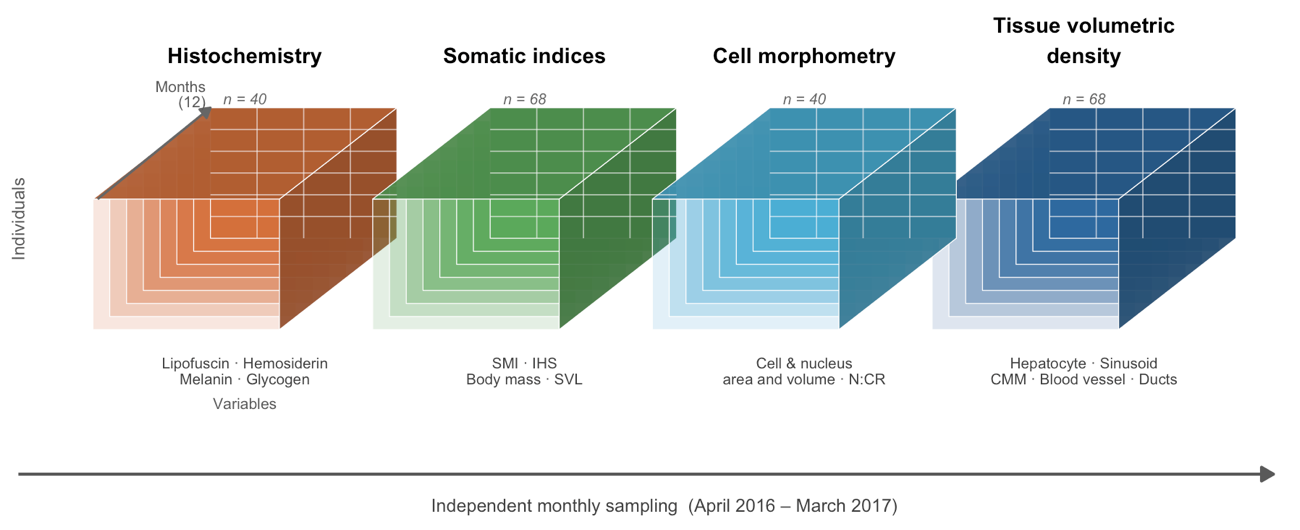

**Fig. S1.** Structure of the multivariate dataset used in this study. Each block represents a data matrix (one of four phenotypic modules) collected from Dendropsophus minutus adult males: Histochemistry (lipofuscin, hemosiderin, melanin, and glycogen area fractions in hepatic cells), Somatic indices (Scaled Mass Index, hepatosomatic index, body mass, and snout–vent length), Cell morphometry (hepatocyte and nucleus area and volume, nucleus-to-cell ratio), and Tissue volumetric density (proportional area occupied by hepatocytes, sinusoids, melanomacrophage centres, blood vessels, and ducts in photomicrographs). Within each block, rows represent individual frogs (n) and columns represent variables. The depth axis corresponds to monthly time points (n = 12), sampled independently from April 2016 to March 2017.

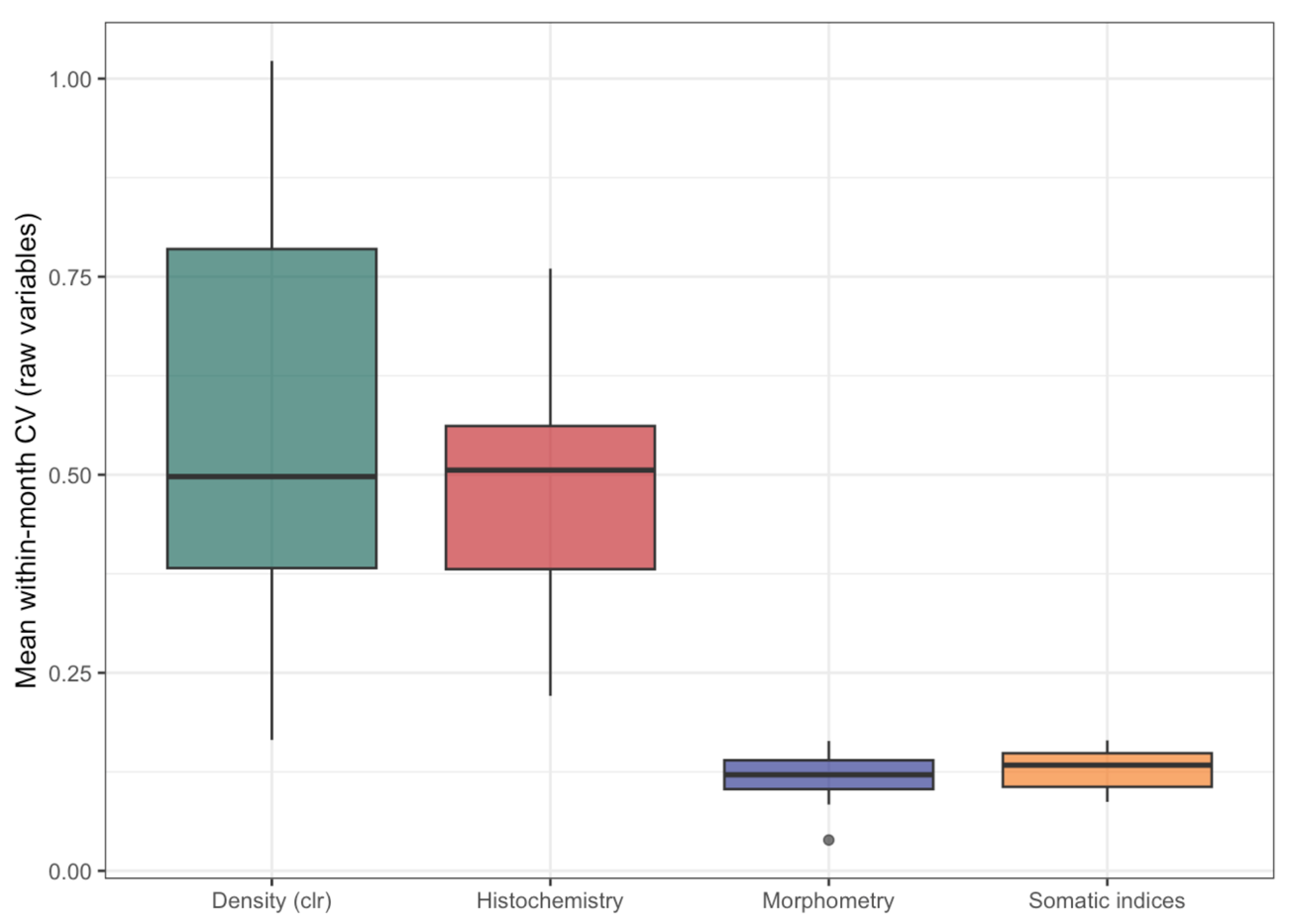

**Fig. S2.** Within-month coefficient of variation (CV) for trajectory speed across the four phenotypic modules. Somatic indices — the fastest module — also show the highest within-month CV, indicating that the speed ranking is not an artifact of differential measurement noise. If noise inflated speed estimates, modules with highest CV would appear fastest spuriously; instead, the pattern is consistent with genuine biological signal driving the observed speed differences.

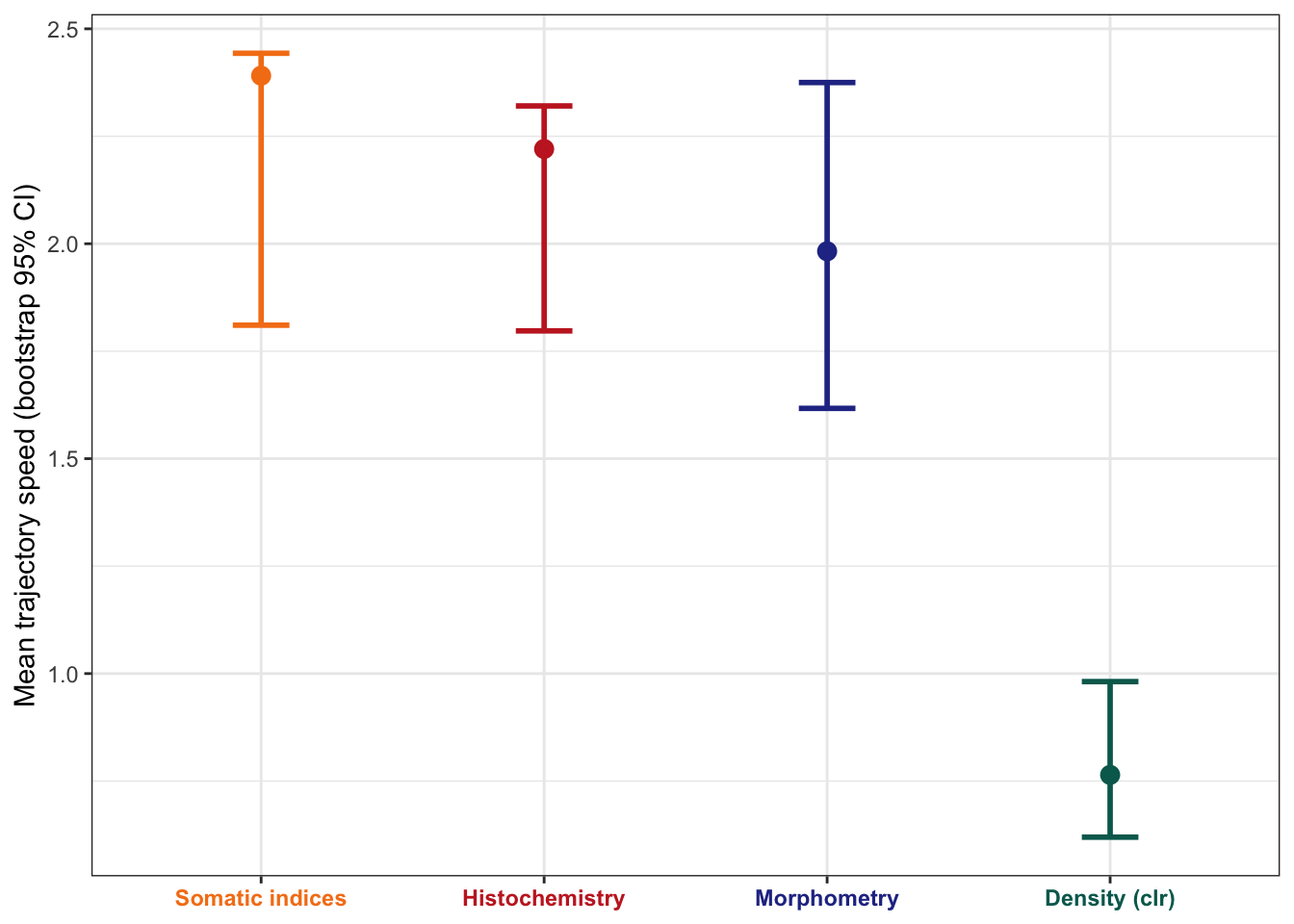

**Fig. S3.** Bootstrap 95% confidence intervals for trajectory speed (mean pairwise distance between consecutive monthly centroids) of each phenotypic module. Speed was computed from standardized between-month distance matrices (999 bootstrap replicates). The fast-responding group (Somatic indices, Histochemistry, Cell morphometry) shows overlapping CIs, while Volumetric density is distinctly slower, supporting the two-tier temporal hierarchy described in the main text.

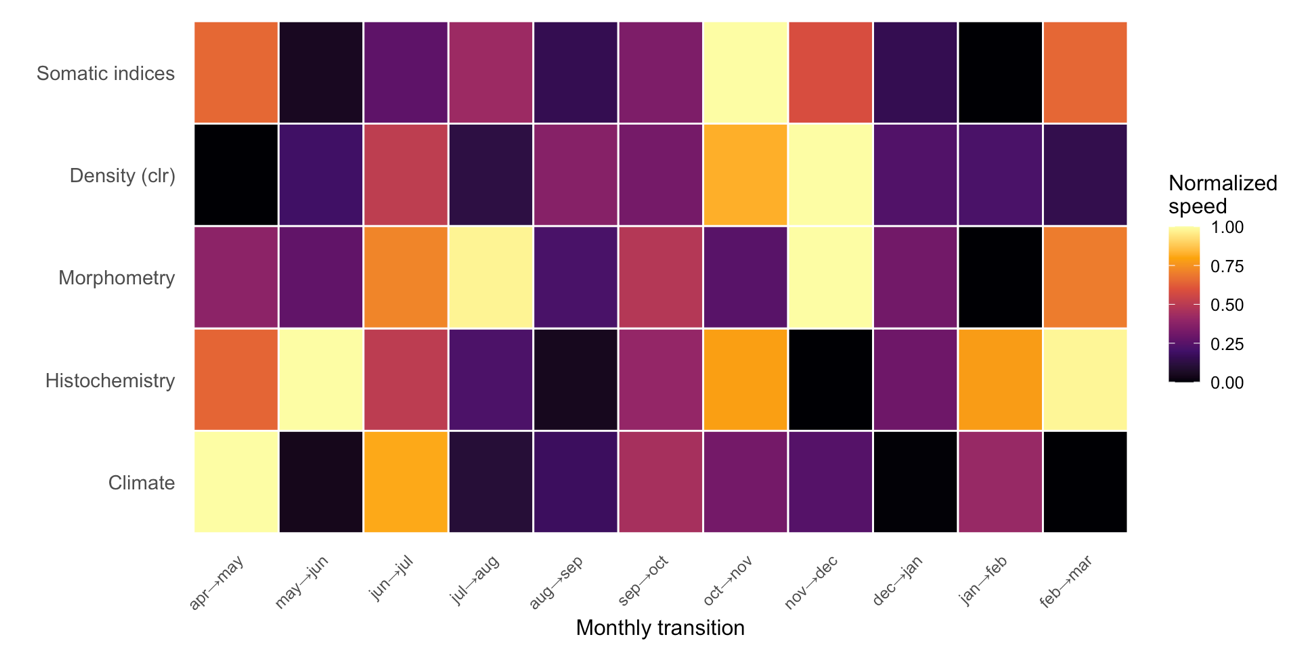

**Fig. S4.** Normalized speed of phenotypic change (displacement in PC1–PC2 space) across consecutive months for each phenotypic module. Warmer colours indicate faster change. Each module reaches its peak rate of change at a different time; these module-specific offsets relative to the shared climatic signal illustrate the temporal response hierarchy described in the main text.

**Table S1.** Full lagged PROTEST results for all edges, both directions, and all lags. P-values are from cyclic-shift permutation (minimum attainable = 1/23 ≈ 0.043).

| **lag** | **m2** | **correlation** | **p_value** | **n_months** | **n_perm** | **edge** | **direction** |
| --- | --- | --- | --- | --- | --- | --- | --- |
| 0 | 0.7909 | 0.4573 | 0.7391 | 12 | 22 | Climate → Histochemistry | forward |
| 1 | 0.7185 | 0.5305 | 0.5217 | 11 | 22 | Climate → Histochemistry | forward |
| 2 | 0.5918 | 0.6389 | 0.1304 | 10 | 22 | Climate → Histochemistry | forward |
| 3 | 0.3789 | 0.7881 | 0.0435 | 9 | 22 | Climate → Histochemistry | forward |
| 0 | 0.7909 | 0.4573 | 0.7391 | 12 | 22 | Climate → Histochemistry | reverse |
| 1 | 0.6777 | 0.5677 | 0.1304 | 11 | 22 | Climate → Histochemistry | reverse |
| 2 | 0.8004 | 0.4468 | 0.6957 | 10 | 22 | Climate → Histochemistry | reverse |
| 3 | 0.7167 | 0.5322 | 0.5217 | 9 | 22 | Climate → Histochemistry | reverse |
| 0 | 0.7706 | 0.4789 | 0.7391 | 12 | 22 | Climate → Morphometry | forward |
| 1 | 0.7502 | 0.4998 | 0.7391 | 11 | 22 | Climate → Morphometry | forward |
| 2 | 0.7269 | 0.5226 | 0.6957 | 10 | 22 | Climate → Morphometry | forward |
| 3 | 0.7354 | 0.5144 | 0.8261 | 9 | 22 | Climate → Morphometry | forward |
| 0 | 0.7706 | 0.4789 | 0.7391 | 12 | 22 | Climate → Morphometry | reverse |
| 1 | 0.6533 | 0.5888 | 0.4348 | 11 | 22 | Climate → Morphometry | reverse |
| 2 | 0.5767 | 0.6506 | 0.3913 | 10 | 22 | Climate → Morphometry | reverse |
| 3 | 0.4217 | 0.7605 | 0.1304 | 9 | 22 | Climate → Morphometry | reverse |
| 0 | 0.3776 | 0.7889 | 0.087 | 12 | 22 | Climate → Density | forward |
| 1 | 0.4166 | 0.7638 | 0.3913 | 11 | 22 | Climate → Density | forward |
| 2 | 0.4205 | 0.7613 | 0.3043 | 10 | 22 | Climate → Density | forward |
| 3 | 0.4587 | 0.7357 | 0.4783 | 9 | 22 | Climate → Density | forward |
| 0 | 0.3776 | 0.7889 | 0.087 | 12 | 22 | Climate → Density | reverse |
| 1 | 0.5375 | 0.6801 | 0.5652 | 11 | 22 | Climate → Density | reverse |
| 2 | 0.7678 | 0.4819 | 0.8696 | 10 | 22 | Climate → Density | reverse |
| 3 | 0.7507 | 0.4993 | 0.8261 | 9 | 22 | Climate → Density | reverse |
| 0 | 0.476 | 0.7239 | 0.0435 | 12 | 22 | Climate → Somatic | forward |
| 1 | 0.547 | 0.6731 | 0.1739 | 11 | 22 | Climate → Somatic | forward |
| 2 | 0.5888 | 0.6412 | 0.3913 | 10 | 22 | Climate → Somatic | forward |
| 3 | 0.7016 | 0.5463 | 0.6522 | 9 | 22 | Climate → Somatic | forward |
| 0 | 0.476 | 0.7239 | 0.0435 | 12 | 22 | Climate → Somatic | reverse |
| 1 | 0.6269 | 0.6108 | 0.4348 | 11 | 22 | Climate → Somatic | reverse |
| 2 | 0.6936 | 0.5535 | 0.6087 | 10 | 22 | Climate → Somatic | reverse |
| 3 | 0.8803 | 0.346 | 0.8696 | 9 | 22 | Climate → Somatic | reverse |
| 0 | 0.8063 | 0.4402 | 0.6087 | 12 | 22 | Histochemistry ↔ Morphometry | forward |
| 1 | 0.6637 | 0.5799 | 0.1304 | 11 | 22 | Histochemistry ↔ Morphometry | forward |
| 2 | 0.814 | 0.4313 | 0.6087 | 10 | 22 | Histochemistry ↔ Morphometry | forward |
| 3 | 0.8297 | 0.4127 | 0.7391 | 9 | 22 | Histochemistry ↔ Morphometry | forward |
| 0 | 0.8063 | 0.4402 | 0.6087 | 12 | 22 | Histochemistry ↔ Morphometry | reverse |
| 1 | 0.7866 | 0.4619 | 0.6087 | 11 | 22 | Histochemistry ↔ Morphometry | reverse |
| 2 | 0.8444 | 0.3945 | 0.8261 | 10 | 22 | Histochemistry ↔ Morphometry | reverse |
| 3 | 0.8797 | 0.3469 | 0.913 | 9 | 22 | Histochemistry ↔ Morphometry | reverse |
| 0 | 0.7549 | 0.4951 | 0.5652 | 12 | 22 | Morphometry → Density | forward |
| 1 | 0.8863 | 0.3373 | 0.9565 | 11 | 22 | Morphometry → Density | forward |
| 2 | 0.8222 | 0.4216 | 0.7826 | 10 | 22 | Morphometry → Density | forward |
| 3 | 0.5182 | 0.6941 | 0.2174 | 9 | 22 | Morphometry → Density | forward |
| 0 | 0.7549 | 0.4951 | 0.5652 | 12 | 22 | Morphometry → Density | reverse |
| 1 | 0.5153 | 0.6962 | 0.2174 | 11 | 22 | Morphometry → Density | reverse |
| 2 | 0.7701 | 0.4794 | 0.5652 | 10 | 22 | Morphometry → Density | reverse |
| 3 | 0.9515 | 0.2202 | 0.9565 | 9 | 22 | Morphometry → Density | reverse |
| 0 | 0.6094 | 0.625 | 0.087 | 12 | 22 | Density → Somatic | forward |
| 1 | 0.4358 | 0.7511 | 0.0435 | 11 | 22 | Density → Somatic | forward |
| 2 | 0.7844 | 0.4643 | 0.5652 | 10 | 22 | Density → Somatic | forward |
| 3 | 0.7785 | 0.4706 | 0.6087 | 9 | 22 | Density → Somatic | forward |
| 0 | 0.6094 | 0.625 | 0.087 | 12 | 22 | Density → Somatic | reverse |
| 1 | 0.5516 | 0.6697 | 0.087 | 11 | 22 | Density → Somatic | reverse |
| 2 | 0.6838 | 0.5623 | 0.4348 | 10 | 22 | Density → Somatic | reverse |
| 3 | 0.8358 | 0.4052 | 0.8261 | 9 | 22 | Density → Somatic | reverse |

**Table S2.** Number of Dendropsophus minutus individuals contributing to each phenotypic module across the 12 sampling months (April 2016 – March 2017). Somatic indices and tissue volumetric density were measured for the full set of 68 frogs; histochemistry and cell morphometry were restricted to a subset of 40 frogs with sufficient histological material for fine-scale measurements (the same individuals contributed to both modules). Months with n ≤ 2 individuals (May, June, August, October, and February for histochemistry and morphometry) are the focus of the sensitivity analyses reported in Figure S3.

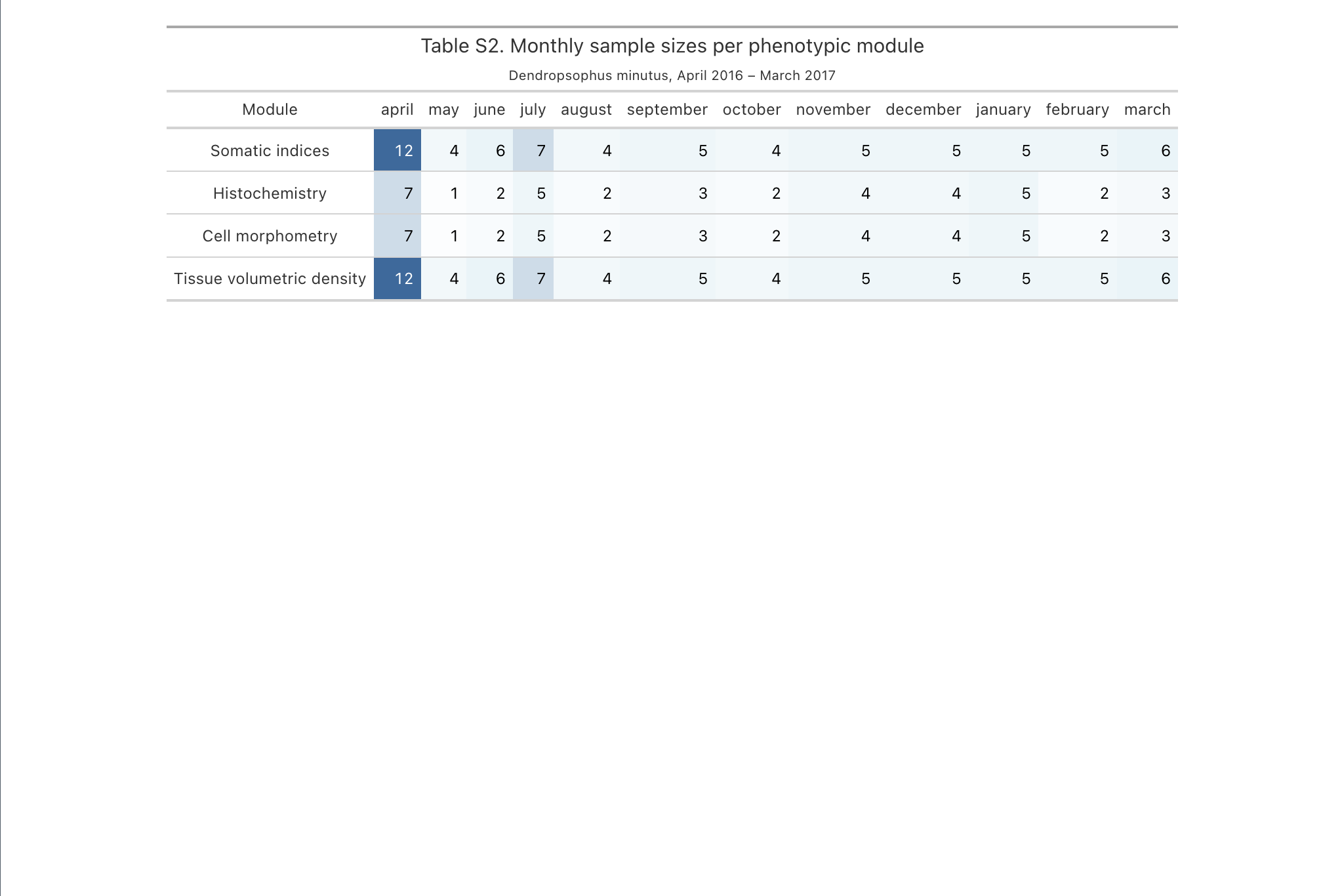

**Table S3.** Trajectory metrics for each phenotypic module: observed value and 95% bootstrap confidence interval (percentile method, 999 replicates resampling individuals within months). Speed is the mean Euclidean displacement per month (ED/month); path length is the total distance traversed over the 12-month cycle (ED); directionality is the ratio of net displacement to total path length (0 = returns to origin, 1 = unidirectional). Modules are ordered by speed.

| **Module** | **Speed (ED/month)** | **Path length (ED)** | **Directionality** |
| --- | --- | --- | --- |
| Somatic indices | 2.39 [1.84, 2.42] | 26.30 [20.25, 26.57] | 0.32 [0.33, 0.43] |
| Histochemistry | 2.22 [1.78, 2.32] | 24.42 [19.63, 25.56] | 0.31 [0.34, 0.41] |
| Cell morphometry | 1.98 [1.65, 2.40] | 21.81 [18.17, 26.36] | 0.38 [0.29, 0.41] |
| Volumetric density (clr) | 0.76 [0.61, 1.00] | 8.41 [6.68, 10.95] | 0.38 [0.28, 0.41] |

**Table S4.** Pairwise differences between phenotypic modules for each trajectory metric, from the paired bootstrap (999 replicates). Δ is the mean difference (module A − module B); the 95% CI is the percentile interval of the bootstrap distribution; P is a two-sided bootstrap probability (2 × the smaller of the proportions of replicates in which the difference was ≤ 0 or ≥ 0). Volumetric density differs from every fast module in both speed and path length (all P < 0.001), whereas the three fast modules do not differ from one another, and no pair differs in directionality.

| **Metric** | **Comparison (A − B)** | **Δ** | **95% CI** | **P** |
| --- | --- | --- | --- | --- |
| Speed | Somatic − Density | 1.343 | [0.984, 1.711] | <0.001 |
| Speed | Histochemistry − Density | 1.275 | [0.928, 1.599] | <0.001 |
| Speed | Morphometry − Density | 1.217 | [0.809, 1.655] | <0.001 |
| Speed | Somatic − Morphometry | 0.127 | [−0.358, 0.563] | 0.559 |
| Speed | Somatic − Histochemistry | 0.069 | [−0.319, 0.454] | 0.745 |
| Speed | Histochemistry − Morphometry | 0.058 | [−0.422, 0.498] | 0.797 |
| Path length | Somatic − Density | 14.776 | [10.823, 18.823] | <0.001 |
| Path length | Histochemistry − Density | 14.021 | [10.204, 17.588] | <0.001 |
| Path length | Morphometry − Density | 13.383 | [8.904, 18.204] | <0.001 |
| Path length | Somatic − Morphometry | 1.393 | [−3.940, 6.195] | 0.559 |
| Path length | Somatic − Histochemistry | 0.755 | [−3.506, 4.998] | 0.745 |
| Path length | Histochemistry − Morphometry | 0.638 | [−4.640, 5.479] | 0.797 |
| Directionality | Somatic − Density | 0.041 | [−0.040, 0.122] | 0.300 |
| Directionality | Somatic − Morphometry | 0.035 | [−0.042, 0.114] | 0.372 |
| Directionality | Histochemistry − Density | 0.032 | [−0.037, 0.104] | 0.392 |
| Directionality | Histochemistry − Morphometry | 0.025 | [−0.039, 0.088] | 0.412 |
| Directionality | Somatic − Histochemistry | 0.009 | [−0.050, 0.069] | 0.741 |
| Directionality | Morphometry − Density | 0.007 | [−0.078, 0.082] | 0.857 |
